# MEIOSIN retains an intrinsic STRA8-independent activity that drives meiotic entry across vertebrate evolution

**DOI:** 10.64898/2026.03.04.709545

**Authors:** Ryuki Shimada, Yukiko Imai, Kimi Araki, Toshihiro Kawasaki, Sakie Iisaka, Shingo Usuki, Kei-ichiro Yasunaga, Sayoko Fujimura, Naoki Tani, Hitoshi Niwa, Shigehiro Kuraku, Noriyoshi Sakai, Kei-ichiro Ishiguro

## Abstract

Meiotic entry in vertebrates has been viewed as a transcriptional switch driven by the MEIOSIN–STRA8 axis, but whether this represents the ancestral regulatory logic of meiosis has remained unclear. Here we combine comparative genomics with genetic and single-cell analyses in zebrafish and mice to show that MEIOSIN retains an intrinsic STRA8-independent activity. We identify Meiosin orthologs in zebrafish and hagfish, vertebrate lineages that lack Stra8, and show that these proteins retain the HMG domain but have lost the bHLH domain required for the canonical MEIOSIN–STRA8 interaction. In zebrafish, *meiosin* is transiently induced at meiotic entry in both sexes, yet its loss selectively disrupts the female germ line, causing failure of oogenesis and female-to-male sex reversal. In mice, MEIOSIN lacking the bHLH domain still initiates key features of meiotic entry and partially activates meiotic target genes, although it fails to support full meiotic progression. Together, these findings identify HMG-containing MEIOSIN as a conserved core regulator of meiotic entry and position STRA8 as a later-acting module that enhances the efficiency and robustness of meiotic gene activation.

## Introduction

Meiosis is the defining developmental transition that enables gamete formation and sexual reproduction. In vertebrates, this transition is triggered by a dedicated transcriptional program that activates meiotic genes as germ cells exit the mitotic cycle and commit to meiotic prophase. In mice, this switch depends on the transient co-expression of Stimulated by Retinoic Acid 8 (STRA8) and Meiosis Initiator (MEIOSIN), which together activate a broad set of meiotic genes in both male and female germ cells (Baltus et al. 2006) (Anderson et al. 2008) (Oulad-Abdelghani et al. 1996) (Menke et al. 2003) (Feng et al. 2021) (Ishiguro et al. 2020). Both proteins contain a basic helix–loop–helix (bHLH) domain and an HMG or HMG-like domain, and together form a central regulatory axis for meiotic entry in mammals (Baltus et al. 2006) (Kojima et al. 2019) (Ishiguro et al. 2020). However, whether this two-factor module represents a universally conserved vertebrate mechanism—or a derived configuration built upon an older core system—has remained unresolved.

This question is particularly important because STRA8 appears to be absent from multiple fish lineages, including zebrafish, despite the continued occurrence of meiosis in these species (Rodriguez-Mari et al. 2013) (Pasquier et al. 2016). Although *Meiosin* orthologs have been reported in several vertebrates, their distribution, domain architecture, and functional dependence on STRA8 remain poorly defined outside mammals. This leaves two major gaps in our understanding: when the MEIOSIN–STRA8 axis emerged during vertebrate evolution, and how meiotic entry is executed in lineages that lack STRA8. Here we address these questions through a vertebrate-wide comparative analysis of *Meiosin* and *Stra8* orthologs, coupled with functional dissection in zebrafish and mice. We show that HMG-containing MEIOSIN exists in zebrafish and hagfish despite the absence of STRA8, that zebrafish *meiosin* is required for female but not overtly for male fertility, and that mouse MEIOSIN retains a STRA8-independent, bHLH-independent capacity to initiate meiotic gene activation. Together, these findings identify MEIOSIN as an evolutionarily conserved core regulator of meiotic entry and position STRA8 as a later reinforcing component of the vertebrate meiotic initiation program. In addition to the shared mechanism in both sexes, meiotic entry in mice is further shaped by sex-specific regulation. STRA8 binds Retinoblastoma family proteins (RB1 and p107) through its LXCXE motif independently of MEIOSIN (Shimada et al. 2023). In female germ cells, this interaction sequesters RB proteins from E2F and derepresses S-phase-associated genes, coupling pre-meiotic DNA replication to meiotic gene activation. Thus, the STRA8–RB and STRA8–MEIOSIN interactions together ensure the timing and fidelity of meiotic entry in oocytes.

MEIOSIN orthologs containing both bHLH and HMG domains have been identified in mammals, birds, reptiles, amphibians, and several fishes (Ishiguro et al. 2020). By contrast, STRA8 orthologs are present in some fish species, such as Southern catfish and Atlantic salmon, but absent from many teleost genomes (Dong et al. 2013) (Li et al. 2016) (Skaftnesmo et al. 2021). In salmon, *stra8* disruption increases germ cell apoptosis, yet spermatogenesis can still proceed, implying the existence of compensatory mechanisms (Skaftnesmo et al. 2021). These observations raise the possibility that the canonical mammalian MEIOSIN–STRA8 module is not the only vertebrate solution for meiotic entry.

## Results

### *Meiosin* orthologs retained by vertebrate species that lack *Stra8*

In mice, MEIOSIN contains both a bHLH domain and an HMG domain (Ishiguro et al. 2020). Our TBLASTN searches using the mouse HMG domain identified MEIOSIN orthologs in coelacanth and other vertebrates, and subsequent cDNA cloning confirmed a MEIOSIN ortholog in cloudy catshark (Fig. S3, see below), that is co-expressed with *Stra8* in the dorsal portion of adult testis (Fig.S1A, B). These data support broad conservation of HMG-containing MEIOSIN across vertebrates. STRA8 is also predicted to contain a bHLH domain and an HMG-like domain (Baltus et al. 2006) (Kojima et al. 2019), and STRA8 orthologs are broadly conserved in vertebrates (Pasquier et al. 2016) (Fig. S3B). However, STRA8 orthologs have not been identified in several fish lineages, including zebrafish and hagfish (Rodriguez-Mari et al. 2013) (Pasquier et al. 2016). We therefore asked whether *Meiosin* orthologs are retained in these lineages despite the apparent loss of *Stra8*.

We identified a potential inshore hagfish *Meiosin* ortholog (ENSEBUG00000016420.1) encoding an HMG-containing protein on the genomic contig FYBX02009704.1 (Yu et al. 2024). Sequence alignment and phylogenetic analysis supported its relationship to jawed vertebrate *Meiosin* genes (Fig. S3C). This hagfish *Meiosin* was specifically expressed in adult testis, similar to *Sycp3*, but not in other examined tissues (Fig.S2A).

We next identified a candidate zebrafish ortholog at the si:ch211-191j22.7 locus on chromosome 21 (Fig.S2B). Because this genomic region contained ambiguous sequence information, we confirmed its transcript sequence by *de novo* assembly of public zebrafish testis RNA-seq data (SRR5378555) followed by RT–PCR and 5′ RACE (Fig. S2B and C). This yielded a full-length cDNA encoding an HMG-containing protein, confirming that the si:ch211-191j22.7 locus corresponds to zebrafish *Meiosin* (Fig. S3A). Southern blotting showed that zebrafish meiosin is present as a single-copy gene despite the teleost genome duplication (Fig.S2D). RT–PCR detected expression specifically in adult testis, but not in other adult tissues examined (Fig.S2E), indicating that zebrafish meiosin is germ cell-enriched, as in mice.

Notably, both zebrafish and hagfish MEIOSIN proteins retain a conserved HMG domain but lack an obvious bHLH domain (Fig. 1A, Fig. S3A). AlphaFold3 predictions likewise identified the HMG domain but not a bHLH domain in either protein (Fig. 1A). By contrast, the predicted three-dimensional structure of the HMG domain was highly conserved among mouse, hagfish, and zebrafish MEIOSIN proteins (Fig. 1B). Thus, MEIOSIN is retained in zebrafish and hagfish, two vertebrate lineages that lack STRA8, and in both cases the protein has lost the bHLH domain while preserving the HMG domain (Fig. 1C). Phylogenetic relationships suggest that bHLH loss occurred independently in these lineages rather than being secondarily gained elsewhere (Fig. 1C). These findings raised two questions: whether zebrafish *Meiosin* still contributes to meiotic entry without a bHLH domain, and how it functions in the absence of STRA8. Together, these findings identify previously unrecognized Meiosin orthologs in zebrafish and hagfish and show that loss of the bHLH domain correlates with the absence of STRA8 in these lineages.

**Figure 1.**
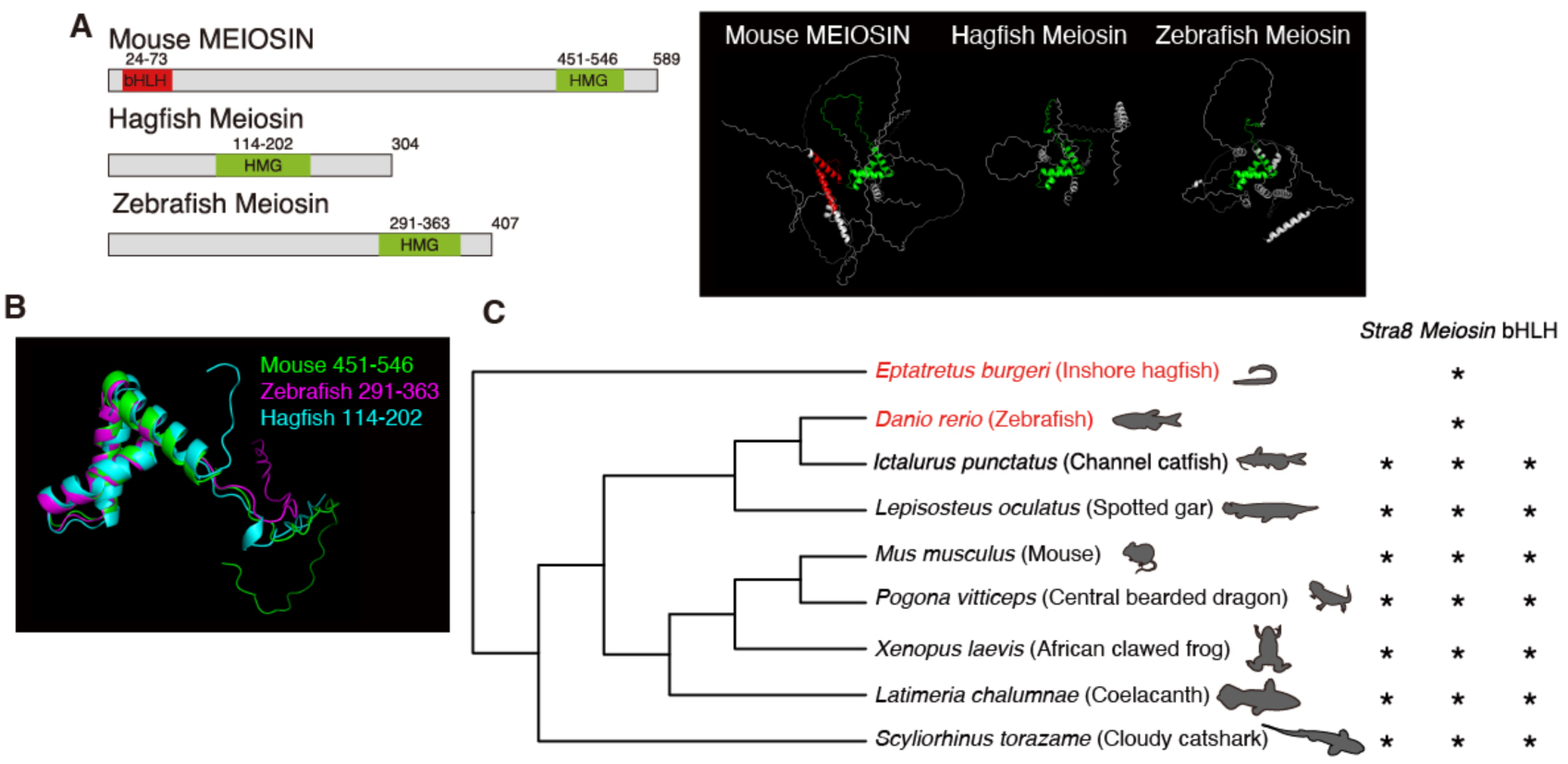
MEIOSIN orthologs identified in zebrafish and hagfish that lack STRA8. **(A)** Domain organization of MEIOSIN orthologs from mouse, hagfish, and zebrafish. Predicted ribbon models generated by AlphaFold3 are shown below. Helices are colored as indicated in the schematics above. **(B)** Superimposed structures of the HMG box from mouse (green), zebrafish (magenta), and hagfish (cyan) MEIOSIN orthologs. Numbers indicate amino acid positions in each species. **(C)** Distribution of *Stra8* and *Meiosin* orthologs mapped onto the vertebrate phylogeny. An asterisk (*) in the bHLH column indicates the presence of a bHLH domain in Meiosin. The phylogeny was generated in R using the Open Tree of Life database (Luna L. Sanchez Reyes 2021).

### Zebrafish *meiosin* marks meiotic entry in both sexes

To define the germ cell populations that express zebrafish *meiosin* (zf *meiosin*), we performed single-cell RNA sequencing (scRNA-seq) on adult WT testes (Fig. 2A, B, C and S4). UMAP analysis resolved 14 transcriptionally distinct germ cell clusters (Fig. 2A, and Fig S4C, Supplementary Data1). Marker expression identified Cluster 8 as spermatogonial stem cells (SSCs) as suggested by the high expression *id4, nanos2* and *dnd1*, Clusters 3/5/9/6 as meiotic prophase cells, and Clusters 2/10/13/7/11 as spermatids (Fig. 2B, C). Based on these markers (Qian et al. 2022; Sposato et al. 2024) (Burgos-Ruiz et al. 2026), we inferred a developmental trajectory from Cluster 8 to Cluster 11.

**Figure 2.**
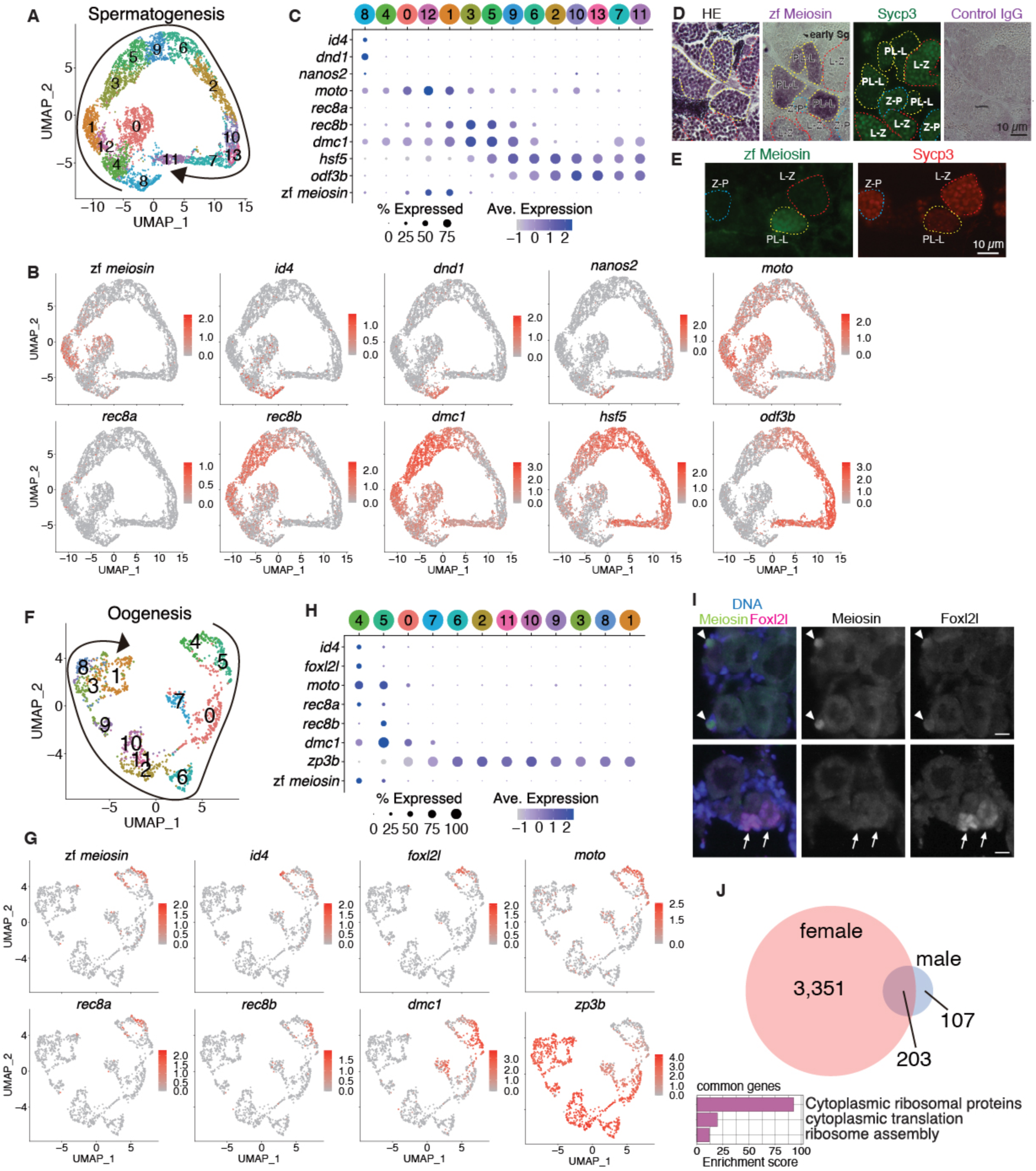
scRNA-seq analysis of male and female germ cells in zebrafish. **(A)** UMAP clustering of scRNA-seq transcriptomes from testicular germ cells isolated from adult WT zebrafish. Arrow indicates the inferred developmental trajectory. See also Fig. S4 for the UMAP of whole testicular cells. Germ cells were separated from somatic cells (encompassing Sertoli cells, Leydig cells, and hemocytes) using established marker genes. **(B)** UMAP plots showing mRNA abundance of meiosin and key marker genes across spermatogenic germ cells. Marker genes: *id4, dnd1* and *nanos2*, spermatogonial stem cells; *moto*, spermatogonia and meiotic-entry cells; *rec8b* and *dmc1*, meiotic prophase spermatocytes; *odf3b*, spermatids. **(C)** Dot plot showing average scaled expression (color gradient) and the fraction of cells with detectable expression (dot size) for key developmental marker genes. **(D)** Serial paraffin sections of adult testis stained with hematoxylin and eosin (HE), anti-Meiosin detected with an AP-conjugated secondary antibody, anti-Sycp3 detected with an Alexa Fluor 488-conjugated secondary antibody, or control rabbit IgG detected with an AP-conjugated secondary antibody. Yellow, red, and cyan dashed lines indicate seminiferous tubules containing preleptotene–leptotene (PL-L), leptotene–zygotene (L-Z), and zygotene–pachytene (Z-P) spermatocysts, respectively. Scale bar, 10 μm. **(E)** Adult testis sections stained for Sycp3 and Meiosin. Dashed lines indicate spermatocysts at the indicated meiotic stages as in (D). Scale bar, 10 μm. **(F)** UMAP clustering of scRNA-seq transcriptomes from ovarian germ cells isolated from *Tg(vas::EGFP)* females at 54 and 85 dpf. Arrow indicates the inferred developmental trajectory. See also Fig. S5 for the UMAP of whole ovarian cells. Germ cells were separated from somatic cells (encompassing Follicle cells, Theca cells, Stromal cells and hemocytes) using established marker genes. **(G)** UMAP plots showing mRNA abundance of meiosin and key marker genes across ovarian germ cells. Marker genes: *id4*, stem cells; *foxl2l*, oocyte progenitors; *moto*, *rec8a*, *rec8b*, and *dmc1*, meiotic-entry/prophase oocytes; *zp3b*, mature oocytes. **(H)** Dot plot showing average scaled expression (color gradient) and the fraction of cells with detectable expression (dot size) for the marker genes shown in (G). **(I)** Serial sections of 42-days post-fertilization (dpf) juvenile ovary stained for Foxl2l (red), Meiosin (green), and DAPI. Arrowheads indicate zf Meiosin-positive cells; arrows indicate strongly Foxl2l-positive cells. Scale bar, 10 μm. **(J)** Overlap between genes enriched in *meiosin*-positive male germ cell clusters (Clusters 12 and 1) and female germ cell clusters (Clusters 4 and 5). Gene ontology enrichment analysis of the shared gene set is shown below. See also Supplementary Data3.

The zebrafish ortholog of mouse *Meioc* (Abby et al. 2016) (Soh et al. 2017) (Mikedis et al. 2023), *moto* (Bowen et al. 2012; Kawasaki et al. 2025), enriched in Clusters 0, 12, and 1, spanning the transition from SSCs to early meiotic prophase. Zf *meiosin* was specifically induced in Clusters 12 and 1, placing its expression in a narrow window at meiotic initiation. Zf *meiosin* was specifically induced in Clusters 12 and 1, placing its expression in a narrow window at meiotic initiation. Consistent with the scRNA-seq data, immunostaining detected Meiosin protein transiently in preleptotene spermatocytes (Fig. 2D, E). Thus, in the male germ line, *meiosin* marks the mitosis-to-meiosis transition.

We next analyzed ovarian germ cells from *Tg(vas::EGFP)* females at 54 and 85 dpf (Fig. 2F, G, H and S5). UMAP analysis identified 12 ovarian germ cell clusters (Fig. 2F, Fig. S5C, and Supplementary Data2). Most clusters corresponded to maturing oocytes, but Cluster 4 expressed the GSC marker *id4* together with the oocyte progenitor marker *foxl2l* (Liu et al. 2022) (Nishimura et al. 2015), suggesting that it contains GSCs and early oocyte progenitors. Cluster 5 appeared to represent a mixed population of late progenitors and early meiotic cells, whereas Clusters 0 and 7 corresponded to meiotic prophase cells (Fig. 2G, H).

Unlike the male germ line, where *rec8b* predominated, both *rec8a* and *rec8b* were expressed in early female clusters, indicating activation before and during early meiotic prophase. Strikingly, *meiosin* was transiently upregulated in Clusters 4 and 5, coincident with *moto* and overlapping with *id4* and *foxl2l*. This suggests that meiotic entry in zebrafish females begins in the GSC/progenitor compartment. Consistent with this interpretation, immunostaining showed that zf Meiosin appears in cells with weak Foxl2l expression, but not in strongly Foxl2l-positive cells (Fig. 2I), placing zf Meiosin induction at the transition out of the progenitor state.

Genes enriched in zf *meiosin*-positive clusters from both sexes were associated with ribosome biogenesis and translation (Fig. 2J, Supplementary Data3). Because failure of meiotic progression in zebrafish oogenesis causes female-to-male sex reversal (Takemoto et al. 2020) (Blokhina et al. 2021) (Imai et al. 2021), the early female-specific transcriptional state accompanying meiosin expression may help couple meiotic entry to commitment to the female pathway.

Thus, zebrafish meiosin is transiently induced at the mitosis-to-meiosis transition in both sexes, with female expression initiating at an unusually early stage in the GSC/progenitor compartment.

### Zebrafish Meiosin is essential for the female germ line

To test the function of zebrafish *meiosin*, we generated a CRISPR–Cas9 frameshift allele by targeting exon 4 and isolating an 11-bp net insertion mutant (Fig. 3A, B). Immunostaining confirmed complete loss of zf Meiosin signal in homozygous mutants, hereafter referred to as *meiosin* KO (Fig. 3C).

**Figure 3.**
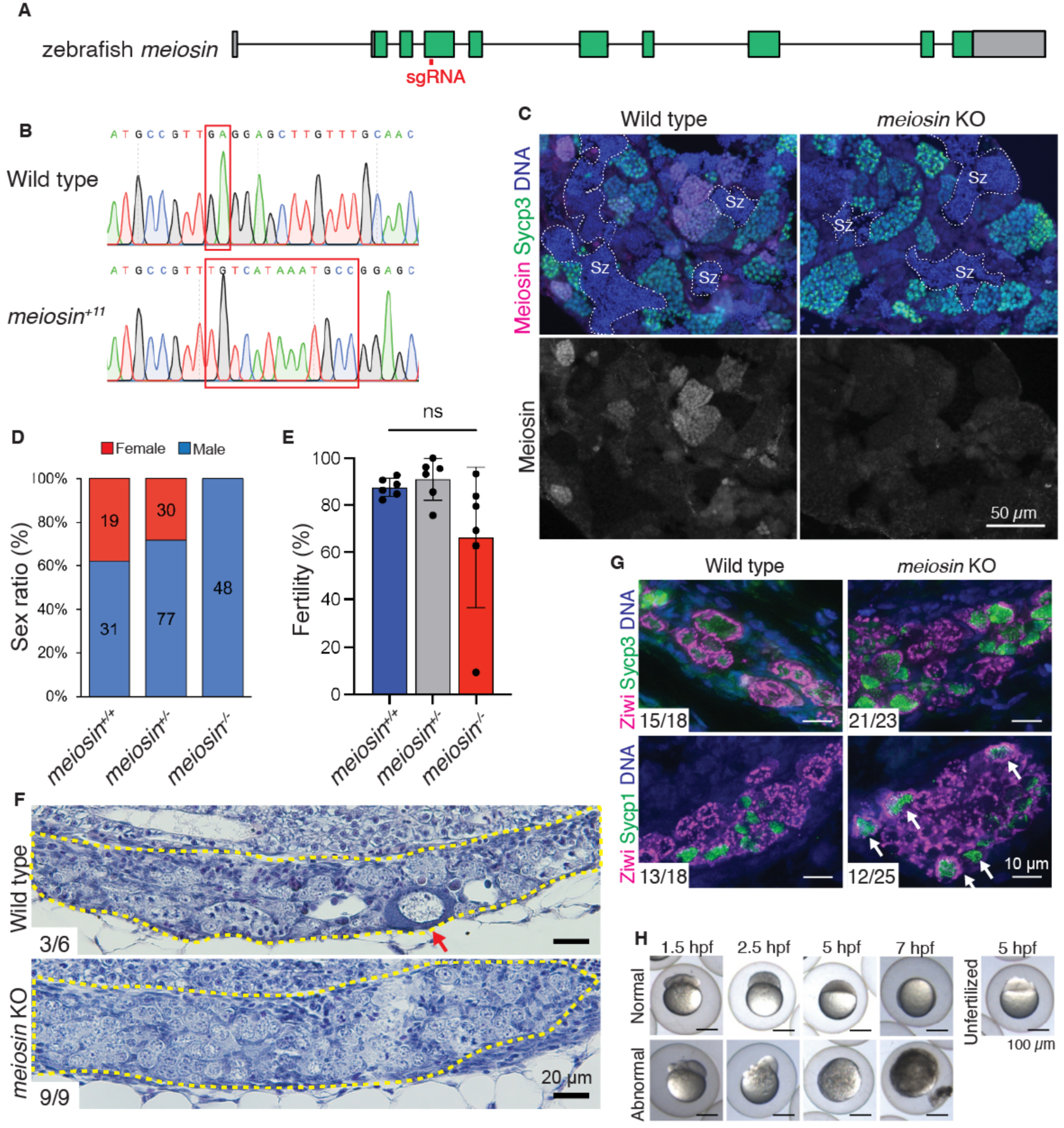
Phenotypes of *meiosin* knockout zebrafish. **(A)** Structure of the zebrafish *meiosin* locus. The protein-coding sequence is shown in green. The sgRNA target site is indicated by a red bar. **(B)** Frameshift mutation allele of *meiosin* generated by CRISPR–Cas9. A 2-bp region was replaced by a 13-bp insertion, resulting in a net insertion of 11-bp (*meiosin^+11^*). **(C)** Adult testis sections from WT and *meiosin* KO fish stained for Sycp3 (green), zf Meiosin (magenta), and DAPI (blue). Sz, spermatozoa. Scale bar, 50 μm. **(D)** Sex ratios of offspring derived from intercrosses of *meiosin* heterozygotes. **(E)** Fertility of WT (*meiosin^+/+^*), heterozygous (*meiosin*^+/-^), and knockout (*meiosin*^-/-^) males. Each male was paired with a WT female. **(F)** HE-stained sections of juvenile gonads from WT and *meiosin* KO zebrafish at 35 dpf. Gonads are outlined by yellow dashed lines. Red arrows indicate stage IB oocytes larger than 20 μm in diameter. Numbers represent the proportion of individuals exhibiting the phenotypes shown. Scale bar, 20 μm. **(G)** Juvenile gonad sections from WT and *meiosin* KO fish stained for Sycp3 or Sycp1 (green), together with Ziwi (magenta) and DAPI (blue). White arrows indicate *meiosin* KO oocytes showing synaptonemal complex formation. Numbers represent the proportion of individuals positive for Sycp3 or Sycp1. Scale bar, 10 μm. **(H)** Embryos derived from paired matings between *meiosin* heterozygous females and WT males are shown. Representative embryos exhibiting normal (upper panels) and abnormal (lower panels) development at each time point are presented. Normal embryogenesis at 1.5, 2.5, 5, and 7 hours post-fertilization (hpf) corresponds to the 8-cell, 256-cell, 30% epiboly, and 75% epiboly stages, respectively. Abnormal development is distinguishable from unfertilized embryos (rightmost panel). Scale bars, 100 µm.

All *meiosin* KO offspring derived from heterozygous intercrosses developed as males, whereas WT and heterozygous siblings developed as both sexes (Fig. 3D). This fully penetrant sex-reversal phenotype indicates that *meiosin* is required for female development. By contrast, *meiosin* KO males produced spermatozoa and displayed fertility comparable to WT males under standard laboratory conditions (Fig. 3E), indicating that Meiosin is not overtly required for male fertility in this setting.

Histological analysis revealed no obvious abnormality in adult males. In juvenile gonads, however, stage IB oocytes were absent in *meiosin* KO animals at 28–35 dpf (Fig. 3C, F and G), indicating arrest before the follicular stage, when oocytes normally enter and remain in dictyate arrest (Selman et al. 1993). Notably, a subset of Sycp3-positive oocytes showed Sycp1-positive synaptonemal complex formation (Fig. 3G), suggesting that mutant oocytes retain a limited capacity to enter and progress through early meiotic prophase I, although they fail to develop further. In addition, many early embryos derived from *meiosin* heterozygous females exhibited abnormal embryogenesis associated with unequal cleavage (Fig. 3H), suggesting dosage sensitivity in the female germ line.

These data show that zebrafish Meiosin is dispensable for male fertility under standard conditions but is crucial for female germ cell development and maintenance of the female developmental pathway.

### The MEIOSIN–STRA8 interaction depends on bHLH domains

The zebrafish phenotype suggested that MEIOSIN can function in the absence of STRA8 and even without a bHLH domain. To define the role of the bHLH domains in the canonical mammalian system, we examined the interaction between mouse MEIOSIN and STRA8. Mouse MEIOSIN contains an N-terminal bHLH domain and a C-terminal HMG domain (Ishiguro et al. 2020), whereas STRA8 contains a putative bHLH domain (Baltus et al. 2006), a glutamate-rich region, an HMG-like domain, and an LXCXE motif required for RB binding (Fig. S7A) (Kojima et al. 2019) (Shimada et al. 2023).

Because bHLH domains frequently mediate protein–protein interactions, and because STRA8 nuclear localization depends in part on MEIOSIN in preleptotene spermatocytes (Ishiguro et al. 2020), we tested whether the two proteins interact through their bHLH regions. In 293T cells, anti-FLAG immunoprecipitation of STRA8-3×FLAG-HA recovered WT MEIOSIN, but not MEIOSIN lacking the bHLH domain (Fig. S6A). Conversely, STRA8 lacking the bHLH domain failed to recover MEIOSIN. By contrast, deletion of the MEIOSIN HMG domain did not abolish the interaction. These results identify the bHLH domains as the principal determinants of the MEIOSIN–STRA8 interaction.

To define the physiological significance of this domain, we generated a *Meiosin*-ΔbHLH mouse line lacking the MEIOSIN bHLH region (Fig. S6B). Meiosin-ΔbHLH males developed smaller testes than controls and lacked post-meiotic spermatids and sperm, closely resembling Meiosin KO testes (Fig. S6C). Female mutants showed ovarian degeneration and a marked reduction in mature follicles, again resembling Meiosin KO animals (Fig. S6D). Thus, the MEIOSIN bHLH domain is required for normal meiosis in both sexes.

*In vivo*, STRA8 localization in postnatal day 17 (PD17) *Meiosin*-ΔbHLH testes was shifted toward the cytoplasm, although some nuclear signal remained, unlike the near-complete loss of nuclear STRA8 in *Meiosin* KO (Fig. 4A). This residual localization contrasts with the interaction defect observed in 293T cells and suggests that in spermatocytes, MEIOSIN may still weakly support STRA8 localization through indirect or context-dependent mechanisms. Together, these results establish the bHLH domains as the principal determinants of the MEIOSIN–STRA8 interaction and demonstrate their requirement for full meiotic function in mice.

**Figure 4.**
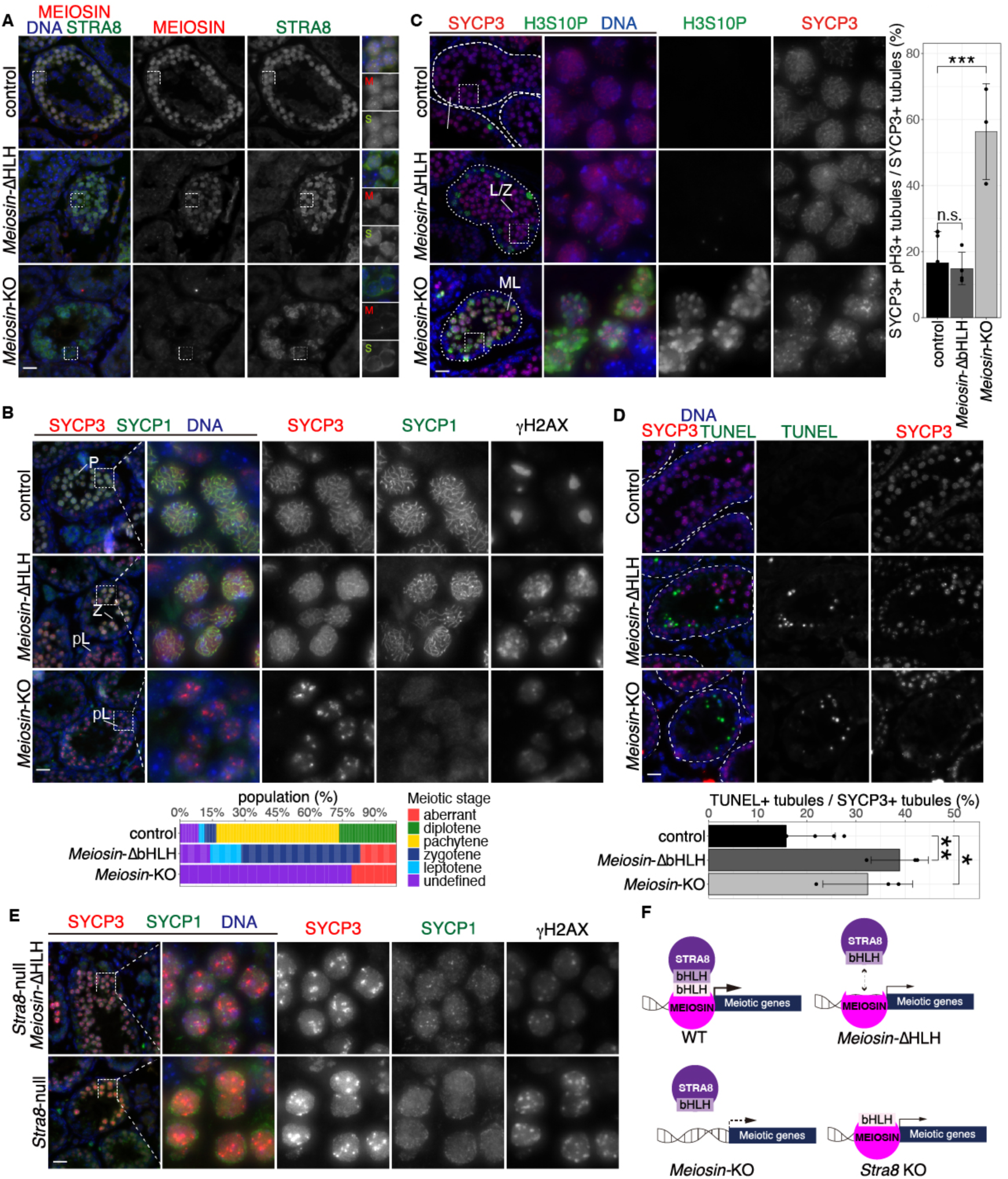
The bHLH domains of STRA8 and MEIOSIN are required for full meiotic progression in mice. **(A)** Seminiferous tubule sections from WT, *Meiosin*-ΔbHLH, and *Meiosin* KO testes at PD17 stained for STRA8, MEIOSIN, and DAPI. Enlarged views are shown on the right. **(B)** Seminiferous tubule sections at PD17 stained as indicated. SYCP3 and SYCP1 mark the synaptonemal complex, and γH2AX marks DNA double-strand breaks. Scale bar, 20 μm. Quantification of meiotic prophase stages is shown below (see also Fig. S7E). pL, preleptotene; Z, zygotene; P, pachytene. **(C)** Seminiferous tubule sections at PD17 stained as indicated. Enlarged views are shown on the right. ML, mitotic-like H3S10P+/SYCP3+ cells; L/Z, leptotene/zygotene. Quantification of seminiferous tubules containing SYCP3+/pH3+ cells among SYCP3+ tubules is shown on the right for control (n = 6), *Meiosin*-ΔbHLH (n = 4), and *Meiosin* KO (n = 3) testes (mean ± s.d.). n.s., not significant (*p* = 0.947); *** *p* = 0.000185. Scale bar, 20 μm. **(D)** Seminiferous tubule sections at PD17 subjected to TUNEL assay together with SYCP3 immunostaining. Scale bar, 20 μm. Quantification of TUNEL-positive tubules among SYCP3-positive tubules is shown below. **(E)** Seminiferous tubule sections at PD17 from WT, *Meiosin*-ΔbHLH/*Stra8*-null, and *Stra8*-null testes stained as indicated. Scale bar, 20 μm. **(F)** Model summarizing meiotic gene activation in WT, *Meiosin*-ΔbHLH, *Meiosin* KO, and *Stra8* KO germ cells.

### MEIOSIN can trigger meiotic entry without its bHLH domain

Despite the gross similarity between *Meiosin*-ΔbHLH and *Meiosin* KO gonads, closer examination revealed clear cellular differences. In *Meiosin*-ΔbHLH testes, at least a subset of SYCP3-positive cells progressed to a more advanced zygotene-like state, as judged by SYCP1-positive synaptonemal complex formation and broad γH2AX staining (Fig. 4B and S6E). These cells did not progress beyond pachytene stage, but they clearly advanced further than *Meiosin* KO cells.

Consistent with our previous work, *Meiosin* KO spermatocytes failed to undergo the mitosis-to-meiosis transition and instead accumulated as mitotic-like cells marked by pH3 and diffuse, patchy SYCP3 staining (Fig. 4C) (Ishiguro et al. 2020). In contrast, such mitotic-like cells were not readily detected in *Meiosin*-ΔbHLH testes, indicating that the transition into the meiotic state had occurred. However, both *Meiosin*-ΔbHLH and *Meiosin* KO testes contained abundant TUNEL-positive tubules (Fig. 4D), showing that germ cells entering prophase are subsequently eliminated by apoptosis.

A similar partial meiotic phenotype has been reported in *Stra8* KO testes (Mark et al. 2008) (Ishiguro et al. 2020). To test whether the residual activity of *Meiosin*-ΔbHLH depends on STRA8, we analyzed *Meiosin*-ΔbHLH/*Stra8*-null testes. Cells with leptotene/zygotene-like features, including partial SYCP1 staining and broad γH2AX signals, were still detected (Fig. 4E), indicating that *Meiosin*-ΔbHLH can initiate aspects of the meiotic program even in the absence of STRA8. However, the double-mutant phenotype was more severe than that of *Meiosin*-ΔbHLH alone, indicating that STRA8 still enhances the residual activity of bHLH-deficient MEIOSIN.

Therefore, MEIOSIN retains a bHLH-independent capacity to trigger meiotic initiation, whereas the bHLH domain is required to drive the program to completion (Fig. 4F).

### The MEIOSIN bHLH domain is required for full transcriptional reprogramming

To define the molecular basis of this partial phenotype, we performed scRNA-seq on spermatogenic cells from WT, *Meiosin*-ΔbHLH, and *Meiosin* KO testes at postnatal day (PD) 15 (Fig. 5 and S7). UMAP analysis identified 15 transcriptionally distinct clusters spanning spermatogenic development (Fig. 5A, Fig. S7E and Supplementary Data4). Cluster 10 corresponded to SSCs, whereas Clusters 0, 12, and 9 represented transitional populations in which *Meiosin* is transiently induced (Fig. 5B).

**Figure 5.**
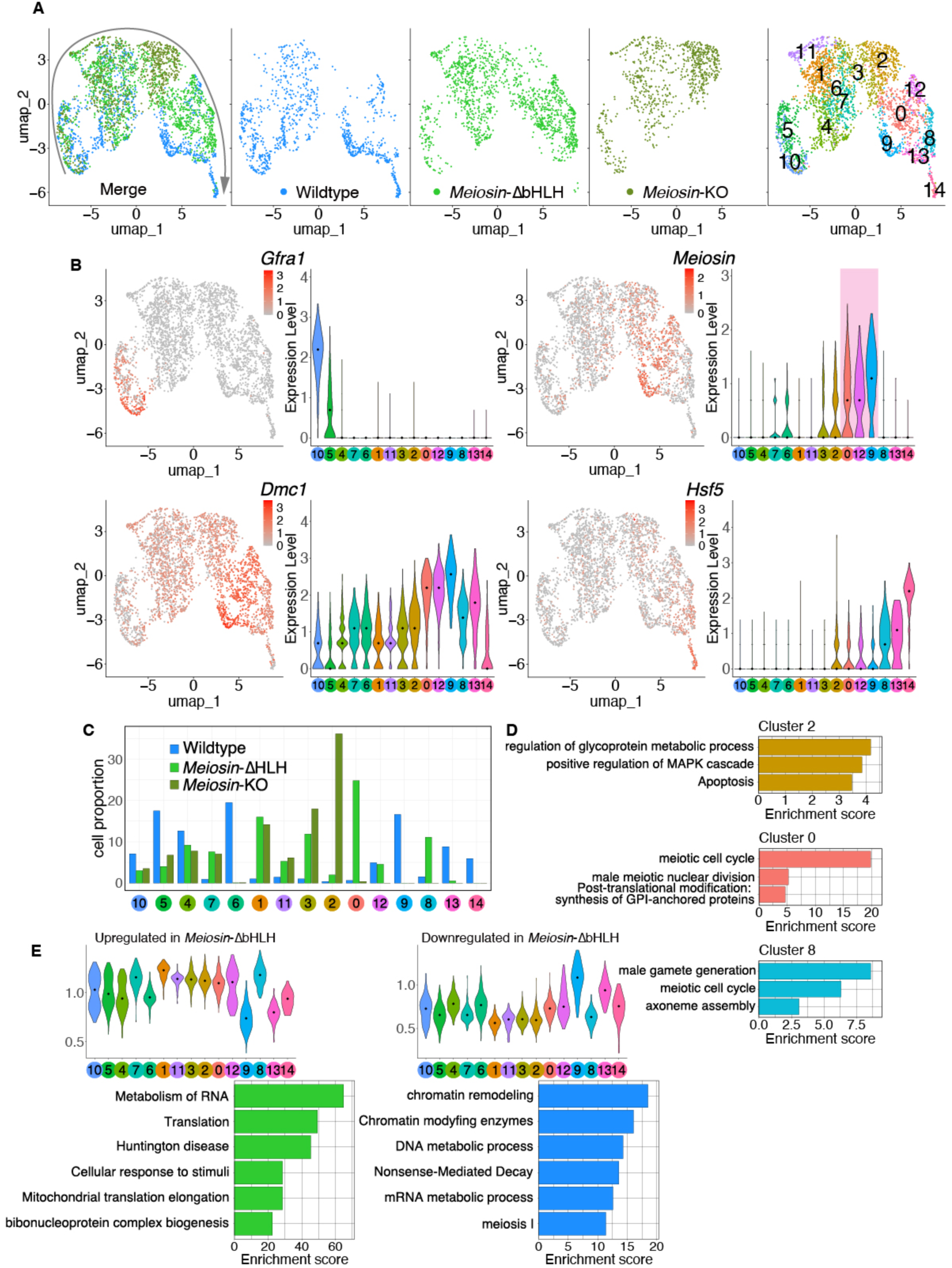
scRNA-seq analysis of WT, *Meiosin*-ΔbHLH, and *Meiosin* KO spermatogenic germ cells. **(A)** UMAP representation of scRNA-seq transcriptomes from spermatogenic germ cells isolated from PD15 WT (Alavattam et al. 2024), *Meiosin*-ΔbHLH, and *Meiosin* KO testes. Cells are colored by genotype (far left) or cluster identity (right). Arrow indicates the inferred differentiation trajectory based on marker gene expression. See also Fig.S7 for the UMAP of whole testicular cells. Germ cells were separated from somatic cells (encompassing Sertoli cells, Leydig cells, and hemocytes) using established marker genes. **(B)** UMAP plots (left) and violin plots (right) showing mRNA abundance of key developmental genes in spermatogenic germ cells. Black dots indicate medians. Marker genes: *Gfra1*, spermatogonial stem cells; *Meiosin*, meiotic initiation (highlighted in pink); *Dmc1*, early meiotic prophase; *Hsf5*, late meiotic prophase. **(C)** Proportion of WT, *Meiosin*-ΔbHLH, and *Meiosin* KO germ cells in each cluster. **(D)** Metascape analysis of the top 100 enriched genes in Clusters 2, 0, and 8, which show genotype-biased cell distributions. Bars indicate enrichment scores (log10(*p-*value)). See also Fig. S7E and Supplementary Data4 for the remaining clusters. **(E)** Violin plots showing average expression of differentially expressed genes in Clusters 0, 9, 12, and 13, comparing *Meiosin*-ΔbHLH and WT spermatocytes. Genes upregulated in *Meiosin*-ΔbHLH are shown on the left, and genes downregulated in *Meiosin*-ΔbHLH are shown on the right, together with their corresponding Metascape analyses. Black dots indicate medians. See also Supplementary Data5.

As expected, *Meiosin* KO cells accumulated in Cluster 2, consistent with failure to initiate meiosis (Fig. 5C). This cluster was enriched for apoptosis-related genes (Fig. 5D). By contrast, *Meiosin*-ΔbHLH cells passed through Clusters 0 and 12, indicating entry into meiosis, but failed to reach late meiotic prophase, as shown by the absence of cells in Cluster 14, where *Hsf5* is strongly induced (Fig. 5B, C). Clusters 0 and 8 were enriched for *Meiosin*-ΔbHLH cells (Fig. 5C) and retained expression of meiotic genes (Fig. 5D), confirming partial activation of the meiotic program. However, this transition was incomplete. Genes normally associated with early spermatogenic growth, particularly those involved in RNA metabolism and translation, remained inappropriately elevated in *Meiosin*-ΔbHLH cells. Conversely, genes involved in chromatin regulation and meiotic prophase progression were reduced relative to WT (Fig. 5E and Supplementary Data5). Thus, *Meiosin*-ΔbHLH cells enter the meiotic state but fail to execute the full transcriptional switch required for orderly meiotic progression.

These findings indicate that the bHLH domain is dispensable for entry into the meiotic state per se, but is required for the full transcriptional reprogramming that supports meiotic progression.

### MEIOSIN activates target genes independently of bHLH, but less efficiently

We next asked how the MEIOSIN bHLH domain contributes to direct target gene activation. Previously defined MEIOSIN-upregulated genes (Ishiguro et al. 2020) were preferentially expressed during meiotic prophase, but showed distinct timing: some, such as *Dmc1*, were induced immediately after *Meiosin* expression began, whereas others, such as *Hsf5*, were induced later, after *Meiosin* expression had already declined (Fig. 5B). We therefore classified these targets into early and late groups (Fig. 6A and B and Supplementary Data6).

**Figure 6.**
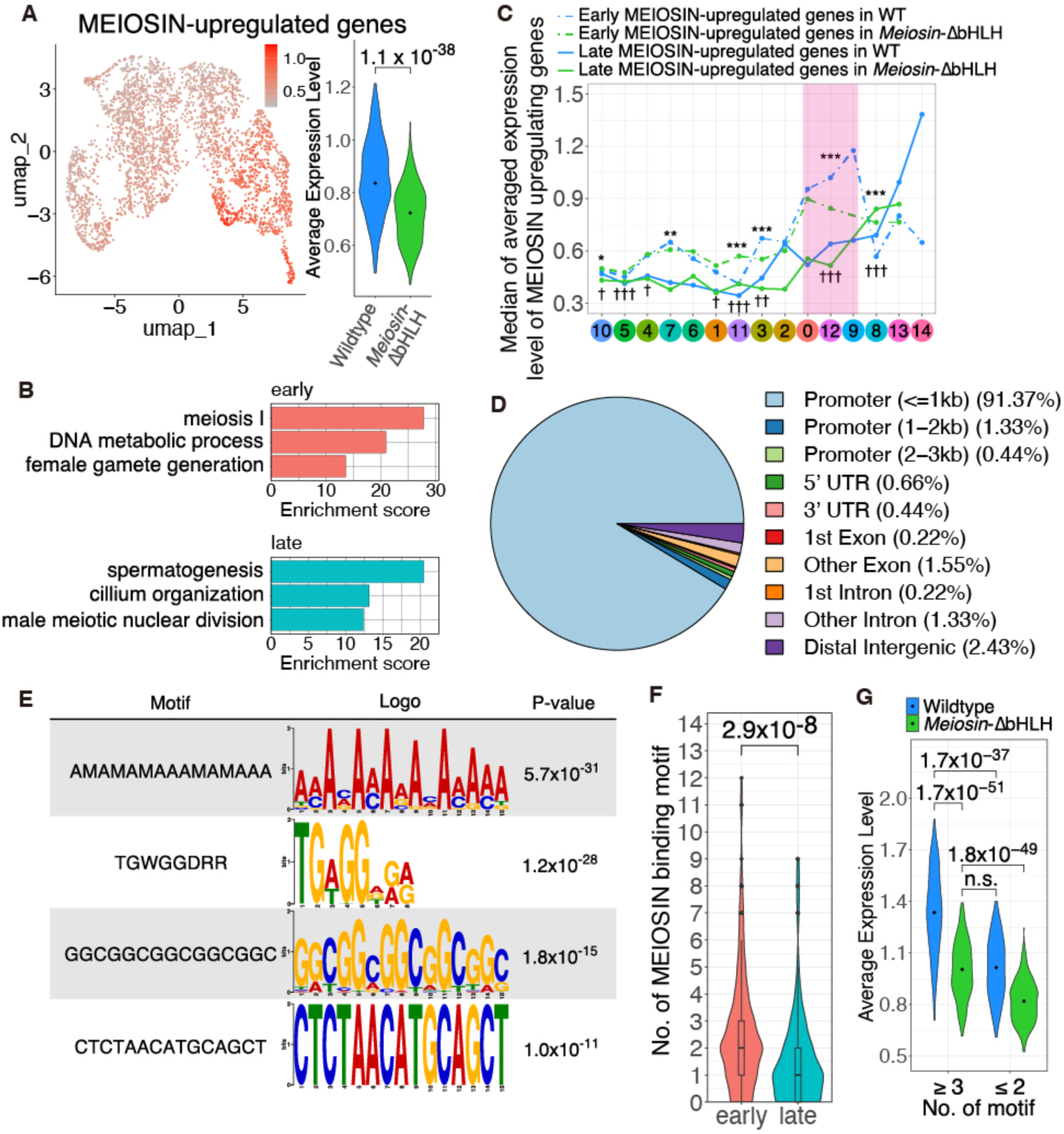
MEIOSIN activates target genes independently of the bHLH domain, but less efficiently. **(A)** UMAP plots showing average mRNA abundance of MEIOSIN-upregulated genes (left) and violin plots showing their average expression in Clusters 0, 12, 9, 8, and 13 (right). Black dots indicate medians. *p*-values were calculated using a two-sided Wilcoxon rank-sum test. **(B)** Metascape analysis of early and late MEIOSIN-upregulated genes. See also Supplementary Data6. **(C)** Median average expression of early and late MEIOSIN-upregulated genes in each cluster for *Meiosin*-ΔbHLH and WT cells. Statistical significance is indicated as *p*-values (early genes: *P < 0.05, ** *p* < 0.01, *** *p* < 0.001; late genes: †*p* < 0.05, ††*p* < 0.01, †††*p* < 0.001). Pink shading indicates the period of *Meiosin* expression. **(D)** Genomic distribution of MEIOSIN-bound peaks associated with MEIOSIN-upregulated genes. **(E)** DNA sequence motifs enriched in MEIOSIN-bound peaks associated with MEIOSIN-upregulated genes. Gray-shaded motifs were excluded from subsequent analyses. **(F)** Violin plot showing the number of MEIOSIN-binding motifs per gene among MEIOSIN-upregulated genes. *p*-values were calculated using a two-sided Wilcoxon rank-sum test. **(G)** Average expression of MEIOSIN-upregulated genes classified by the number of MEIOSIN-binding motifs per gene. *p*-values were calculated using a two-sided Wilcoxon rank-sum test.

Meiosin-ΔbHLH cells still activated both groups of MEIOSIN targets, but early genes were induced less strongly than in WT cells (Fig. 6C). Late genes appeared broadly preserved in Cluster 0/12/9 (Fig. 6C), although their strongest induction normally occurred in Cluster 14, which was absent in *Meiosin*-ΔbHLH testes (Fig. 5C). These results indicate that the bHLH domain is not absolutely required for transcriptional activation by MEIOSIN, but markedly enhances the amplitude and fidelity of target gene induction.

To examine how this residual activity is encoded, we reanalyzed our previously published MEIOSIN ChIP–seq data (Ishiguro et al. 2020) and found that 91.4% of MEIOSIN-binding peaks associated with genes upregulated in this study lie near transcription start sites (Fig. 6D). Motif analysis identified two prominent non-repetitive enriched motifs —TGWGGDRR and CTCTAACATGCAGCT— (Fig. 6E). Early MEIOSIN target genes contained more MEIOSIN-binding motifs than late targets, and among early genes, those with three or more motifs were expressed more strongly than those with fewer motifs in both WT and *Meiosin*-ΔbHLH cells (Fig. 6F). Notably, expression of ≥3-motif genes in *Meiosin*-ΔbHLH cells approached that of ≤2-motif genes in WT cells (Fig. 6G) suggesting that multiple binding sites can partially compensate for reduced MEIOSIN activity in the absence of the bHLH domain.

We then compared WT, *Meiosin* KO, and *Stra8*-null testicular scRNA-seq datasets (Fig. S8A–C). While *Meiosin* KO cells accumulated in Cluster 2 and *Stra8*-null cells in Cluster 0—both corresponding to stages prior to meiotic entry (Fig. S8D, E and Supplementary Data7). Although both mutants arrested before full meiotic progression (Fig. S8B), *Stra8*-null cells expressed MEIOSIN-upregulated genes at higher levels than *Meiosin* KO cells (Fig. S8F), indicating that MEIOSIN can activate target genes independently of STRA8. STRA8 therefore acts not as an obligate trigger, but as an enhancer of MEIOSIN-dependent transcription.

To identify factors that might support this STRA8-independent activity, we generated mice carrying both 3xFLAG-HA tags at the C-terminus of MEIOSIN and performed tandem-affinity purification of endogenously tagged MEIOSIN followed by mass spectrometry (Fig. S9A, B). In addition to STRA8, we identified components of the TIP60/NuA4 histone acetyltransferase complex, including TRRAP, RUVBL1, and RUVBL2, as MEIOSIN-associated proteins (Fig. S9C, D, Supplementary Data8). AlphaFold-based predictions further suggested that intrinsically disordered regions of MEIOSIN acquire structure upon complex formation with TRRAP, both in mouse (Fig. S9E, F) and, to a more limited extent, in zebrafish (Fig. S9G and H). These observations support a model in which MEIOSIN has intrinsic transcriptional activity that can be potentiated by chromatin-associated cofactors even in the absence of STRA8.

Taken together, these results show that MEIOSIN can activate meiotic target genes independently of both its bHLH domain and STRA8, but that robust target gene induction requires the reinforced activity of the MEIOSIN–STRA8 axis.

## Discussion

### An ancestral HMG-based core for meiotic entry

Our findings support a two-layer model for the evolution of meiotic entry in vertebrates. In this model, HMG-containing MEIOSIN constitutes an ancestral core regulator capable of initiating the meiotic transcriptional program, whereas STRA8 and the bHLH-mediated MEIOSIN–STRA8 interaction represent later-added modules that reinforce, amplify, and stabilize this core activity. This framework explains why meiosis can still be initiated in species that lack STRA8 and why bHLH-deficient MEIOSIN retains partial function in mice.

The evolutionary logic of this model is further supported by domain architecture. MEIOSIN proteins in zebrafish and hagfish retain a conserved HMG domain and broadly disordered regions, but lack the bHLH domain (Fig.1, S3A). Given that MEIOSIN orthologs in other vertebrate lineages that retain STRA8 also retain the bHLH domain, the most parsimonious interpretation is that the ancestral vertebrate MEIOSIN protein possessed both domains, and that in the zebrafish and hagfish lineages, *Stra8* was lost secondarily during evolution. Under this scenario, once STRA8 was lost, the bHLH domain of MEIOSIN—whose principal role is to mediate the canonical MEIOSIN–STRA8 interaction—would have become dispensable and was subsequently lost independently in these lineages. Thus, the loss of the MEIOSIN bHLH domain in zebrafish and hagfish is likely not a primitive state, but a secondary simplification following the loss of STRA8.

The fission yeast meiotic regulator Ste11 (Sugimoto et al. 1991) shows a similar combination of an HMG domain and extended non-globular regions, suggesting that HMG-based control of meiotic entry may reflect a deeply conserved regulatory solution. Our study therefore identifies the HMG domain as the minimal conserved architectural feature required for MEIOSIN function and places the bHLH-dependent interaction with STRA8 as a derived enhancement rather than the ancestral core. We propose that HMG-containing MEIOSIN represents an evolutionarily conserved core module for meiotic entry, whereas STRA8 and the bHLH-mediated MEIOSIN–STRA8 interaction act as lineage-specific amplifiers of this ancestral program.

### STRA8 strengthens, but does not define, meiotic initiation

Our data refine the functional relationship between MEIOSIN and STRA8. In the prevailing mammalian framework, the two factors are often viewed as a single obligate module. However, the present results indicate that MEIOSIN retains intrinsic activity in the absence of both STRA8 and its own bHLH domain (Figure 4-6). This residual activity is sufficient to trigger aspects of meiotic entry and to induce a subset of target genes, especially when multiple MEIOSIN-binding motifs are present near promoters.

The combined *Meiosin*-ΔbHLH; *Stra8*-null mutant exhibited phenotypes similar to those of the Stra8-null mutant alone (Figure 4E). This suggests that, even in the absence of the bHLH domain, MEIOSIN function is enhanced by STRA8. Although the interaction between MEIOSIN and STRA8, mediated by the bHLH domain, may be required for formation of a stable complex (Figure S6A), the two proteins may still interact weakly in a bHLH-independent manner. This raises the possibility that, in fish species lacking Stra8, such as zebrafish and hagfish, MEIOSIN function may be enhanced by an as-yet-unidentified factor through a bHLH-independent mechanism. Thus, STRA8 is not the sole trigger of meiotic entry; rather, it enhances the efficiency, magnitude, and completeness of MEIOSIN-dependent transcription.

This interpretation helps reconcile previous genetic observations. In some mouse strain backgrounds, *Stra8* mutants can still progress into early meiotic prophase (Mark et al. 2008) (Ishiguro et al. 2020), consistent with the idea that MEIOSIN alone can partially activate the meiotic program, leading to phenotypic variation among different mouse strains. Such phenotypic variation may reflect differences in chromatin state, cofactor availability, or promoter architecture that modulate the threshold for MEIOSIN function.

Our data further suggest that MEIOSIN regulates transcription through both direct and indirect mechanisms. We found that MEIOSIN target genes can be categorized into early and late groups based on their expression dynamics (Figure 6). Notably, late genes remain upregulated even after Meiosin expression has ceased, indicating that at least part of the downstream transcriptional program is maintained indirectly, likely through secondary regulatory factors induced by MEIOSIN.

The strong enrichment of NuA4/TIP60 components among MEIOSIN-associated proteins (Figure S9) further suggests that chromatin remodeling and histone acetylation help determine the competence of germ cells to respond to MEIOSIN.

The germ cell specificity of both MEIOSIN and STRA8 function is also consistent with this view. Even if MEIOSIN has intrinsic transcriptional activity, it likely operates only in a permissive chromatin environment established in germ cells at the onset of meiotic differentiation. A key next step will therefore be to define the epigenetic features that license MEIOSIN to activate meiotic genes and to understand how STRA8 and other cofactors increase the robustness of this response.

This framework explains how meiotic entry can still occur in the absence of STRA8, while also accounting for the enhanced efficiency and robustness conferred by the MEIOSIN–STRA8 complex in species that retain both factors.

### A reduced meiotic-entry module in zebrafish

The zebrafish data reveal a striking sexual asymmetry. Although *meiosin* is induced at meiotic entry in both sexes (Figure 2), its loss selectively disrupts the female germ line (Figure 3), causing arrest of oogenesis and complete female-to-male sex reversal, while males remain fertile under standard conditions. This indicates that in zebrafish, the female developmental program is highly dependent on timely *meiosin* function, specifically for meiotic entry, whereas the male germ line either tolerates reduced meiotic-entry activity or engages compensatory mechanisms.

One possibility is that the female pathway imposes a narrower developmental window during which meiotic entry must be coupled to commitment to oogenesis. In this context, reduced or delayed activation of meiotic genes may trigger failure of the female program and redirection toward male development. By contrast, the male germ line may be buffered by broader developmental timing, redundant regulators, or a chromatin environment that more readily supports residual MEIOSIN activity. The finding that *meiosin* is induced in the GSC/progenitor compartment in females (Figure 2G, I), earlier than in males, is consistent with the idea that meiotic entry is more tightly integrated with lineage commitment in the female germ line.

Our mouse genetic dissection suggests how such a reduced system could operate. Because MEIOSIN can activate target genes independently of STRA8 and without a bHLH domain (Figure 6), zebrafish MEIOSIN likely relies on alternative cofactors and/or promoter configurations to achieve sufficient activity. Identifying those cofactors—particularly proteins that may stabilize MEIOSIN or recruit chromatin-modifying complexes in a STRA8-independent manner—will be essential for understanding how meiotic entry is implemented in lineages that have lost the canonical MEIOSIN–STRA8 axis.

Defining the cofactors and chromatin states that potentiate zebrafish MEIOSIN will be key to understanding how a reduced, STRA8-independent meiotic entry module operates in vertebrate lineages that have lost the canonical MEIOSIN–STRA8 axis.

## Methods

### Animals

#### Mice

*Δ*HLH mutant *Stra8^Δ^*^HLH^-*3xFLAG-HA-p2A-GFP* knock-in (*Stra8 ^Δ^*^HLH^-*3FH* KI), *Meiosin Δ*HLH mutant knock-in (*Meiosin ^Δ^*^HLH^-KI) mice and other knockout and knock-in mice were congenic with the C57BL/6 background (age: postnatal day 17, 4- weeks and 8-weeks old). Wild type *Stra8-3xFLAG-HA-p2A-GFP* knock-in (*Stra8wt-3FH* KI), *Stra8-null GFP* knock-in (*Stra8 null GFP*-KI), *Meiosin* KO mice (age: postnatal day 17, 4-weeks and 8-weeks old) were generated as described in our previous studies (Ishiguro et al. 2020) (Shimada et al. 2023). Male mice were used for, histological analysis of testes, immunostaining of testes and sc-RNA-seq experiments (age: postnatal day 15, and 8-weeks old). Female mice were used for histological analysis of the ovaries (age: 8-weeks old). Whenever possible, each knockout animal was compared to littermates or age-matched non-littermates from the same colony, unless otherwise described. Animal experiments were approved by the Institutional Animal Care and Use Committee (Kumamoto University approval F28-078, A2022-001, A2024-028, Chiba University approval A8-110).

#### Fish

Male and female hugfish were purchased from Shinshomaru (Enoshima, Japan). Zebrafish (*Danio rerio*) were maintained under standard conditions as described in The Zebrafish Book (Westerfield 1995). The wild-type AB line (see ZFIN at http://zfin.org/action/genotype/view/ZDB-GENO-980210-28) was used to generate the *meiosin-like* KO zebrafish line by the CRISPR-Cas9 mutagenesis based on published protocols (Chen et al. 2017) (Hwang et al. 2013). Template DNA for single-guide RNA (sgRNA) synthesis was prepared by amplification with a primer specific to *meiosin-like* exon 4: 5′ - taatacgactcactataGGTGCAAACAAGCTCCTCAAgttttagagctagaa -3′, and a universal reverse primer: 5′-aaaagcaccgactcggtgccactttttcaagttgataacggactagccttattttaacttgctatttctagctctaaaac -3′, and T4 DNA polymerase. After purification of the template DNA, sgRNA was transcribed in vitro with a MEGAscript T7 kit (Ambion) and purified with a MEGA clean-up kit (Ambion). Wild-type AB embryos were injected at the 1- or 2-cell stage with 2.3 nl of a mixture of 10 pmol/μl Cas9 NLS protein (abm) and 100 ng/μl *meiosin* sgRNA. Founders were backcrossed with AB fish, and the F1 siblings were screened by genotyping using a specific primer pair; Meiosin P4F: 5′-acagactaaagactgcgtgagat -3′ and Meiosin P4R: 5′- tgacgagttgctttgtctgttg -3′. The mutations were verified by Sanger sequencing with vector-specific primers M13 Fwd (-20): 5′-GTAAAACGACGGCCAG -3′ and M13 Rev: 5′- CAGGAAACAGCTATGAC -3′, after cloning the PCR products into the pCRII vector using TOPO TA cloning kit (Invitrogen), Heterozygous *meiosin-like* knockout fish carrying a 11-bp frameshift insertion in exon 4 was intercrossed to obtain homozygote mutant fish.

### Generation of *Meiosin*-Δ HLH mutant *Meiosin*^Δ^ ^HLH^ knock-in mouse and genotyping

Δ HLH mutant *Meiosin* knock-in (*Meiosin*^Δ^ ^HLH^ KI) mouse was generated by introducing Cas9 protein (317-08441; NIPPON GENE, Toyama, Japan), tracrRNA (GE-002; FASMAC, Kanagawa, Japan), synthetic crRNA (FASMAC) and ssODN into C57BL/6N fertilized eggs using electroporation. The synthetic crRNAs were designed to direct GAGAACAATTACCTGTCAGT (agg) of the *Meiosin* Exon 2, TCATGTCAATGTTTCTCTGG (agg) of the *Meiosin* Exon 4, ssODN: 5′-TTGCCATTGCTGCTGTGGAGCTTGAGCAAGGCCTTGGTCATGTCAATGTTCCTGTCAGTAGGACTTGGAGATCTAGCTCTGTGCTGATCAGAGAAACATG -3′ was used as a homologous recombination template. The electroporation solutions contained 10μM of tracrRNA, 10μM of synthetic crRNA, 0.1 μg/μl of Cas9 protein, ssODN (1μg/μl) in Opti-MEM I Reduced Serum Medium (31985062; Thermo Fisher Scientific). Electroporation was carried out using the Super Electroporator NEPA 21 (NEPA GENE, Chiba, Japan) on Glass Microslides with round wire electrodes, 1.0 mm gap (45-0104; BTX, Holliston, MA). Four steps of square pulses were applied (1, three times of 3 mS poring pulses with 97 mS intervals at 30 V; 2, three times of 3 mS polarity-changed poring pulses with 97 mS intervals at 30 V; 3, five times of 50 mS transfer pulses with 50 mS intervals at 4 V with 40% decay of voltage per each pulse; 4, five times of 50 mS polarity-changed transfer pulses with 50 mS intervals at 4 V with 40% decay of voltage per each pulse).

The *Meiosin*^Δ^ ^HLH^ KI allele (341 bp) was identified by PCR using the following primers; Gm4969-13926F : 5′- GTCCTATTTTAGGAACTCTAGGTGC -3′ and Meiosin delHLH-1R: 5′- CAGGAAGGGGAAGGTAGCCAATTGG -3′. The wild-type allele (212 bp) was identified by PCR using the following primers; Gm4969-13926F and Gm4969-14136R: 5′-GTAGAGAACAATTACCTGTCAGTAGG -3′. The PCR amplicons were verified by sequencing. For avoiding potential mosaicism of the mutant allele, the F0 mutant founders were backcrossed with C57BL/6N to segregate *Meiosin*^Δ^ ^HLH^ KI allele and establish heterozygous lines. Since homozygous mouse with *Meiosin*^Δ^ ^HLH^ KI allele was infertile, female and male mice were maintained as *Meiosin*^Δ^ ^HLH^ KI heterozygous mutant. *Meiosin*^Δ^ ^HLH^ KI allele encodes MEIOSIN lacking 10aa-70aa.

### Generation of *Meiosin*-3xFLAG-HA mutant knock-in mouse and genotyping

3xFLAG-HA was inserted by introducing Cas9 protein (317-08441; NIPPON GENE, Toyama, Japan), tracrRNA (GE-002; FASMAC, Kanagawa, Japan), synthetic crRNA (FASMAC) and ssODN into C57BL/N fertilized eggs using electroporation. The synthetic crRNAs were designed to direct CACCCCTAACTGCCAGTGAG (AGG) of the *Meiosin* Exon 14, ssODN: 5′-ATGCTAAGCAGGTGGCAGAGCCCAGCAGGACTGCCCgTCTCATTAGGCGTAGTCGGGCACGTCGTAGGGGTAtccCTTGTCATCGTCATCCTTGTAATCGATGTCATGATCTTTATAATCACCGTCATGGTCTTTGTAGTCtccCTGGCAGTTAGGGGTGGGCAGTGAGGTGAGGCCCAAAGACAC -3′ was used as a homologous recombination template. The electroporation solutions contained 10μM of tracrRNA, 10μM of synthetic crRNA, 0.1 μg/μl of Cas9 protein, ssODN (1μg/μl) in Opti-MEM I Reduced Serum Medium (31985062; Thermo Fisher Scientific). Electroporation was carried out using the Super Electroporator NEPA 21 (NEPA GENE, Chiba, Japan) on Glass Microslides with round wire electrodes, 1.0 mm gap (45-0104; BTX, Holliston, MA). Four steps of square pulses were applied (1, three times of 3 mS poring pulses with 97 mS intervals at 30 V; 2, three times of 3 mS polarity-changed poring pulses with 97 mS intervals at 30 V; 3, five times of 50 mS transfer pulses with 50 mS intervals at 4 V with 40% decay of voltage per each pulse; 4, five times of 50 mS polarity-changed transfer pulses with 50 mS intervals at 4 V with 40% decay of voltage per each pulse).

Using following primers, wildtype allele (647bp) and the *Meiosin*-3FH KI allele (732bp) were identified by PCR; Gm4969-29318F: 5′- CACTAAGGCACGCAGGTTCAGCCGC -3′ and Gm4969-29964R: 5′- GAGGTAAAGGGTTTGAGTTCAGACC -3′.

### Transfection and cell culture

293T cells were cultured in DMEM, supplemented with 10% FBS, 2 mM L-glutamine, 100 U/mL penicillin and 100 μg/mL streptomycin at 37 °C with 5% CO_2_. For STRA8-MEIOSIN interaction analyses, over expression of the Dox-inducible trans- gene, 3.6 x 10^6^ 293T cells were transfected with the expression vectors and pCAG-PBase using Lipofectamine 2000 followed by culture for 2 days. All plasmids used in this study are as following; pBRPBCAG-STRA8-3FH-IN (WT STRA8-3xFLAG-HA)

pBRPBCAG-STRA8-3FH dHLH-IN (dHLH STRA8-3xFLAG-HA) encoding STRA8 lacking 17aa-84aa.

pBRPBCAG-Gm4969-IN (WT MEIOSIN)

pBRPBCAG-Gm4969 dHMG-IN (dHMG MEIOSIN) encoding MEIOSIN lacking 10aa-70aa.

pBRPBCAG-Gm4969 dHLH-IN (dHLH MEIOSIN) encoding MEIOSIN lacking 484aa-534aa..

### Preparation of cell extracts and immunoprecipitation

Chromatin-bound and -unbound extracts were prepared as described previously (Tani et al. 2022). Briefly, testicular cells were suspended in low salt extraction buffer (20 mM Tris-HCl pH 7.5, 100 mM KCl, 0.4 mM EDTA, 0.1% TritonX100, 10% glycerol, 1 mM β-mercaptoethanol) supplemented with Complete Protease Inhibitor (Roche). After homogenization, the soluble chromatin-unbound fraction was separated after centrifugation at 100,000*g* for 30 min. The chromatin bound fraction was extracted from the insoluble pellet by high salt extraction buffer (20 mM HEPES-KOH pH 7.0, 400 mM KCl, 5 mM MgCl_2_, 0.1% Tween20, 10% glycerol, 1 mM β-mercaptoethanol) supplemented with Complete Protease Inhibitor. The solubilized chromatin fraction was collected after centrifugation at 100,000*g* for 30 min at 4°C.

Immuno-affinity purification was performed with anti-FLAG M2 monoclonal antibody-coupled magnetic beads (Sigma-Aldrich) from the testis chromatin-bound fractions of *Meiosin*-3xFLAG-HA-p2A-GFP knock-in mice. For negative control, mock immuno-affinity purification was done from the testis chromatin-bound and -unbound fractions from the age-matched wild type mice. The beads were washed with high salt extraction buffer for chromatin-bound proteins and low salt extraction buffer for chromatin-unbound proteins. The anti-FLAG-bound proteins were eluted by 3xFLAG peptide (Sigma-Aldrich). The second immuno-affinity purification was performed anti-HA 5D8 monoclonal antibody-coupled Magnet agarose (MBL M132-10). The bead-bound proteins were eluted with 40 µl of elution buffer (100 mM Glycine-HCl [pH 2.5], 150 mM NaCl), and then neutralized with 4 µl of 1 M Tris-HCl [pH 8.0].

The 3xFLAG-HA tagged version of WT STRA8, the STRA8^ΔHLH^ and STRA8^ΔE^ proteins were immunoprecipitated with anti-FLAG M2 monoclonal antibody-coupled magnetic beads (Sigma-Aldrich) from 293T cell extracts expressing *Stra8* ^wt^-*3FH*, *Stra8* ^ΔHLH^-*3FH* and STRA8^ΔE^ -*3FH* respectively. The immunoprecipitated proteins were run on SDS-PAGE and immunoblotted. Immunoblot was conducted for the detection of TRRAP in a Tris-glycine buffer containing 20% methanol, and 0.05% SDS. Immunoblot image was developed using ECL prime (GE healthcare) and captured by FUSION Solo (VILBER).

### Mass spectrometry

Mass spectrometry was conducted as previously decribed (Tani et al. 2022). Briefly, the immunoprecipitated proteins were run on 4-12% NuPAGE gel (Thermo Fisher Scientific) by 1cm from the well and stained with SimplyBlue (Thermo Fisher Scientific) for in-gel digestion. The gel containing the proteins was excised and cut into pieces approximately 1mm size. The proteins in these gel pieces were then reduced with DTT (Thermo Fisher Scientific), alkylated with iodoacetamide (Thermo Fisher scientific), and digested overnight at 37°C using trypsin and Lysyl endopeptidase (Promega) in a buffer containing 40mM ammonium bicarbonate, pH 8.0. The resulting peptides were analyzed using an Advance UHPLC system (ABRME1ichrom Bioscience) connected to a Q Exactive mass spectrometer (Thermo Fisher Scientific), with the raw mass spectra processed using Xcalibur (Thermo Fisher Scientific). The raw LC-MS/MS data were analysed against the SwissProt protein/translated nucleotide database, restricted to *Mus musculus* using Proteome Discoverer v 1.4 (Thermo Fisher Scientific) with the Mascot search engine version2.5 (Mascot Science). A decoy database comprising either randomised or reversed sequences from the target database was used to estimate false discovery rate (FDR), and the Percolator algorithm was used to evaluate false positives. The search results were filtered against a global FDR of 1% to achieve a high level of confidence. Proteins that detected in the control sample were removed from further analysis.

Proteins identified by LC–MS/MS are shown after excluding those also detected in anti-FLAG/ anti-HA immunoprecipitates from wild-type testis extracts. The proteins identified by more than two distinct peptide hits are listed together with their Swiss-Prot accession numbers, gene symbols, amino acid lengths, Mascot scores, and numbers of peptide hits.

### Antibodies

The following antibodies were used for immunoblot (IB) and immunofluorescence (IF) studies in mice: rabbit anti-H3S10P (IF, 1:2000, Abcam: ab5176), rabbit anti-SYCP1 (IF, 1:1000, Abcam ab15090), rabbit anti-TRRAP (WB, 1:1000, Abcam ab73546), rat anti-SYCP3 and guinea pig anti-SYCP3, rabbit and rat anti-STRA8, rabbit and guinea pig anti-MEIOSIN N-terminal (a.a. 1-224) and rabbit and guinea pig anti-MEIOSIN C-terminal (a.a. 405-589) as described previously (Ishiguro et al. 2020). Rabbit anti-zf Vasa (IF, 1:200, GeneTex GTX128306) was used for cloudy catshark. For zebrafish samples, the following antibodies were used: Rabbit anti-zf Meiosin (IF, 1:300, WB, 1:1000, this paper), Mouse anti-zf Foxl2l (IF, 1:100, this paper). Rabbit anti-Ziwi (IF, 1:500) (Houwing et al. 2007) (Kawasaki et al. 2025), guinea pig anti-zf Sycp3 (IF, 1:200) (Imai et al. 2021), rabbit anti-zf Sycp3 (IF,1:200) (Saito et al. 2011), and rat anti-zf Sycp1 (IF, 1:200) (Saito et al. 2014).

### Histological analysis

For hematoxylin and eosin staining, mouse testes, epididymis and ovaries were fixed in Bouin solution, and embedded in paraffin. Sections were prepared on CREST-coated slides (Matsunami) at 6 μm thickness. Cloudy catshark and zebrafish testes, and zebrafish juvenile bodies were fixed in 4% paraformaldehyde for overnight at 4°C or Bouin solution, and embedded in paraffin. Sections were prepared on CREST slides (Matsunami) at 4 µm. The slides were deparaffinized and stained with hematoxylin and eosin.

### Immunofluorescence microscopy of mouse testis and ovary

For immunofluorescence staining, the testes were fixed in 4% PFA for several hours at 4℃ and submerged sequentially in 10%, 20% and 30% sucrose at 4℃, then the fixed ovaries were embedded in Tissue-Tek O.C.T. compound (Sakura Finetek) and frozen at -80℃. Cryosections were prepared on the MAS-GP typeA-coated slides (Matsunami) at 6 μm thickness, and then air-dried. The testes were fixed in 4% paraformaldehyde in PBS at pH 7.4. The sections were blocked for 30 min in 3% BSA containing 0.1% TritonX100 in TBS (TBS-T), and incubated at 4℃ with the primary antibodies in a blocking solution. After three washes in TBS-T, the sections were incubated for 90 min at room temperature with Alexa-dye-conjugated secondary antibodies (1:1000; Invitrogen) in TBS-T. DNA was counterstained with Vectashield mounting medium containing DAPI (Vector Laboratory). TUNEL assay was performed using MEBSTAIN Apoptosis TUNEL Kit Direct (MBL 8445). DNA was counterstained with Vectashield mounting medium containing DAPI (Vector Laboratory). Statistical analyses, and production of graphs and plots were done using R.

### Immunofluorescence microscopy of fish testis and juvenile body

Cloudy catshark and zebrafish testes, and zebrafish juvenile bodies, were fixed in 4% paraformaldehyde overnight at 4 °C, embedded in paraffin, and sectioned at 4 µm on CREST slides (Matsunami). Sections were deparaffinized in xylene and rehydrated through a graded ethanol series (100%, 90%, 80%, 70%, and 50%), followed by four washes in PBS-0.1% Tween-20 for 5 min each. Antigen retrieval was performed in 1× Immuno Saver (Nisshin EM) at 98 °C for 45 min, after which sections were washed once in PBS-0.5% Triton X-100 for 15 min and three times in PBS-0.1% Tween-20 for 5 min each. After blocking with 2% BSA-PBS for 30 min at room temperature, sections were incubated with primary antibodies diluted in the blocking solution or Can Get Signal Immunostain Solution (Toyobo) overnight at room temperature. Following three washes in PBS-0.1% Tween-20, sections were incubated with Alexa dye–conjugated secondary antibodies (1:200 for anti-zf Sycp1 and anti-zf Sycp3; 1:500 for anti-zf Meiosin and anti-zf Foxl2l) for 2 h at room temperature. After three washes in PBS-0.1% Tween-20, sections were counterstained with 0.5 µg/mL DAPI for 15 min and mounted in ProLong Glass (Invitrogen).

### Imaging

Immunostaining images were captured with DeltaVision (GE Healthcare). The projection of the images was processed with the SoftWorx software program (GE Healthcare). All images shown were Z-stacked. For counting seminiferous tubules and embryonic ovaries, immunostaining images were captured with BIOREVO BZ-X710 (KEYENCE), and processed with BZ-H3A program. XY-stitching capture by 40x objective lens was performed for multiple-point color images using DeltaVision. Images were merged over the field using SoftWorx software program. Bright field images were captured with OLYMPUS BX53 fluorescence microscope and processed with CellSens standard program.

### Antibody production

Polyclonal antibodies specific for zebrafish Sycp2 was described in the previous study (Takemoto et al. 2020). Polyclonal antibodies against zebrafish Meiosin (a.a. 1-407) were generated by immunizing rabbits. Polyclonal antibody against zebrafish Foxl2l (a.a. 1-260) was generated by immunizing mouse. His-tagged recombinant proteins were produced by inserting cDNA fragments in-frame with pET19b (Novagen) in *E. coli* strain BL21-CodonPlus (DE3)-RIPL (Agilent), solubilized in a denaturing buffer (6 M HCl-Guanidine, 20 mM Tris-HCl pH 7.5) and purified by Ni-NTA (QIAGEN) under denaturing conditions. The antibodies were affinity-purified from the immunized serum with immobilized antigen peptides on CNBr-activated Sepharose (GE healthcare).

### 5′-Full RACE

5′ Full RACE was performed using 5′-Full RACE Core Kit (TaKaRa Bio). Total RNA was isolated from the adult testis of zebrafish using TRIzol (Thermo Fisher Scientific). First cDNA was synthesized using Superscript III from total RNA with a 5′ end-phosphorylated zf *Meiosin* primer, zfMeiosin-5′-RACE-pR; 5′- AGTTGGAGTCTGTAG -3′. The hybrid DNA-RNA was degraded by treatment with RNase H at 30°C for 1 hour. After ethanol precipitation, the single stranded cDNA was ligated using T4 RNA ligase. The ligated product was amplified using PrimeSTAR MAX (TaKaRa) with following primers; zfMeiosin_S1: 5′-ACAGACAAAGCAACTCGTCAC -3′ and zfMeiosin_A1: 5′-AGTCTGTGAGTCCCTCTTATTG -3′. Amplified product was further amplified using MightyAmp (TaKaRa) with following primers; zfMeiosin_S2: 5′-GTTGAAGACAGTGAAGCATCAC -3′ and zfMeiosin_A2: 5′-TGCGACTGTAAGTGTTCATG -3′. PCR product was cloned into pCR2.1 by TA cloning using a TOPO TA Cloning Kit (Thermo Fisher Scientific). and sequenced to determine the sequence.

### Southern blot

10 μg of zebrafish genomic DNA was digested by restriction enzymes and run on 0.8% agarose gel electrophoresis followed by transfer to Hybond N+ membrane (GE) in 0.4N NaOH. The membrane was subjected to hybridization in QuickHyb (Agilent #201220) with ^32^P-labeled probe that encompasses genomic region of meiosin-like Exon2- Exon5 at 65°C. Autoradiography image was acquired by the Typhoon FLA7000 Biomolecular imager (Cytiva).

### Single-cell RNA-sequencing

Testes were collected from wild type zebrafish at adulthood. Ovaries were collected from *Tg(vas::EGFP)* females at 54dph. The ovaries were also collected from *Tg(vas::EGFP, gata6SAGFF254A, UAS::NTR*), in which nitroreductase are expressed in oocytes larger than 40µm. Metronidazole was treated days before dissection and the ovaries were collected from females at 85dph.

Single-cell suspensions were prepared by incubating testes and ovaries with Accutase (Innovative Cell Technologies, Inc.) for 5min at 37°C. After incubation, DMEM with 10% FBS was added to block Accutase and aggregate was disrupted by pipetting. Then, cell suspensions were filtered through a 35-μm cell strainer sieve (BD Bioscience). Cells were collected by centrifugation and re-suspended in PBS containing 0.1% BSA, and dead cells and debris were removed using Cell Sorter SH800 (SONY). Collected cells were re-suspended in DMEM containing 10% FBS. Resulting approximate 1000 ∼2000 single-cell suspensions were loaded on Chromium Controller (10X Genomics Inc.). Single cell RNA-seq libraries were generated using Chromium Single Cell 3’ Reagent Kits v3 following manufacturer’s instructions, and sequenced on an Illumina HiSeq X to acquire paired end 150 nt reads. The number of used embryos, the total numbers of single cells captured from ovaries, mean depth of reads per cell, average sequencing saturation (%), the number of detected genes, median UMI counts/cell, total number of cells before and after QC, are shown in Figure S2. Sequencing data are available at DDBJ Sequence Read Archive (DRA) under the accession.

### Statistical analysis of scRNA-seq

Fastq files were processed and aligned to the zebrafish transcriptome (GENCODE vM23/Ensembl 98) using the 10X Genomics Cell Ranger v 4.0 pipeline. Further analyses were conducted on R (ver.4.4.1) (R Core Team, 2019) via RStudio (ver.2024.04.2+764) (RStudio Team, 2018). Quality assessment of scRNA-seq data and primary analyses were conducted using the Seurat package for R (v5.1.0) (Butler et al. 2018) (Stuart et al. 2019). Contaminated somatic cells were excluded, and only the cells that expressed more than 200 genes were used for further analysis to remove the effect of low-quality cells. The scRNA-seq data were merged and normalized using SCTransform function built in Seurat. The dimensional reduction analysis and visualization of cluster were conducted using RunUMAP function built in Seurat. The clustering of cells was conducted using FindNeighbors and FindClusters built in Seurat with default setting. Determination of differentially expressed genes (DEGs) was performed using FindAllMarkers function built in Seurat with following options, only.pos = T, min.pict = 0.25 and logfc.threshold = 0.25, for the identification of marker genes in each cluster, and FindMarker function built in Seurat with default settings to characterize the arbitrary group pf cells. RNA velocity analysis was conducted using the RNA velocyto package for python (Python 3.7.3) and R with default settings (La Manno et al. 2018), and visualized on UMAP plots built in Seurat.

### Gene enrichment analysis

Gene enrichment analyses were performed using Metascape (Zhou et al. 2019) with default settings and the results were visualized using R. To characterize the feature of each cluster, top100 representative genes in each cluster were used.

### Classification of MEIOSIN upregulating genes

Original MEIOSIN upregulating genes were reported in previous report (Ishiguro et al. 2020). The genes that detected in scRNA-seq data were selected for further analysis. The genes that expressed in cluster 0, 12, 9, and 8 (E1) were determined using FindMarker function built in Seurat with following option, only.pos = T. Highly expressed genes in clusters 13 and 14 compared to clusters 0, 12, 9, and 8 were determined using FindMarker function built in Seurat, with a threshold p_val_adj < 0.05 (Late High). Among original MEIOSIN upregulating gene list, genes that also listed in Late High genes were defined as Late genes. Among genes commonly listed in both original MEIOSIN upregulating genes and E1 genes, genes that not listed as Late genes were defined as Early genes.

### PCR with reverse transcription

Total RNA was isolated from hagfish tissues using TRIzol (Thermo Fisher). Hagfish cDNA was generated from total RNA using Superscript III (Thermo Fisher) followed by PCR amplification using Quick-Taq (TOYOBO) and template cDNA. Cloudy catshark and zebrafish cDNAs were generated from total RNAs using PrimeScript RT-PCR kit (Takara) followed by PCR amplification using KOD Plus Neo (Toyobo) or GoTaq (Promega). Amplified PCR products were cloned into the pCRII vector using TOPO TA cloning kit (Invitrogen), and sequenced with M13 Fwd (-20) and M13 Rev primers.

Sequences of primers used for RT-PCR were as follows:

Hagfish Ddx4 _F1: 5′- GAAGTGCGCTCACTGTCAGAG -3′

Hagfish Ddx4_R1: 5′- CATGACCTCCTCTGCCAACAC -3′

Hagfish Sycp3_F1: 5′- ATGGCCTCTGATCAAGATGACATG -3′

Hagfish Sycp3_R1: 5′- TCCTGGTGCAGAGTCTCCAG -3′

Hagfish Gapdha_F: 5′- CAGCAATGCTTCCTGCACTACC -3′

Hagfish Gapdha_R: 5′- CGTTGTCGTACCATGACACCAAC -3′

Hagfish meiosin-like_F1: 5′- CCCAATGAACAGCCAGACCAAG -3′

Hagfish meiosin-like_R1: 5′- CTGTCCCGTACACAGAATCGGTAG -3′

Zf actb1_F1: 5′- GACGACCCAGACATCAGGGAGTG -3′

Zf actb1_R1: 5′- CACGGACAATTTCTCTTTCGGCTG -3′

Zf meiosin-like_F1: 5′- CCAGGGAGAAAGTCATTGGA -3′

Zf meiosin-like_R1: 5′- AGGCCGTACTGGTGTTTTTG -3′

Cloudy shark Tbp_F: 5′-CGGGAACCTCGTACAACTGC-3′

Cloudy shark Tbp_R: 5′-GGCAGCAGGCAAACGAAATG-3′

Cloudy shark Sycp3_F: 5′-CACTGGGGAGCAGCTATGAGA-3′

Cloudy shark Sycp3_R: 5′-GAAAAGCTTCTGTTGCTGCCG-3′

Cloudy shark Stra8_F: 5′-ATGGAGAGCTCTGGAGATTG-3′

Cloudy shark Stra8_R: 5′-TTACAGATCATCATCGAATGTCTC-3′

Cloudy shark Meiosin_F: 5′-ATGGCCTCCATAATTGAGGTGAAGC-3′

Cloudy shark Meiosin_R: 5′-GTCAGGCAGCATCTTACAGTAGATGG-3′

### Fertilization tests and embryogenesis of zebrafish

To assess male fertility, three independent pair matings per genotype were performed, in which three *meiosin^-/-^*, *meiosin^+/-^* and *meiosin^-/-^* sibling males were individually mated with a wild-type female in the same day. Each pair subjected to two repeated matings. Statistical analysis was performed with the Mann-Whitney two-tailed test using GraphPad Prism 11 software. To examine embryogenesis of *meiosin^+/-^* female-derived embryos, embryos obtained from two independent paired crosses of *meiosin^+/-^* and wild-type male were kept at 28°C. Images were captured with a Leica MX16 FA microscope equipped with a Leica DFC310 FX camera.

### Molecular phylogenetic analysis

Protein sequences were collected from the NCBI and Ensembl databases. The deduced amino acid sequences were aligned with the MAFFT v7.505 (Katoh and Standley 2014) using the L-INS-i method. The aligned sequences were trimmed with trimAl v1.4.rev15 (Capella-Gutierrez et al. 2009) to remove unreliably aligned sites using the ‘-gappyout’ option. The maximum-likelihood tree was inferred with iqtree 1.6.1 (Nguyen et al. 2015) using the JTT+I+G4 model for *Meiosin* and JTT+G4 model for *Stra8*, and for evaluating the confidence of the nodes, the ultrafast bootstrap resampling with 1000 replicates was performed.

## Data availability

All data supporting the conclusions are present in the paper and the supplementary materials. Raw sequence data generated in this study were publicly available as of the date of publication. Sequencing data have been deposited in DDBJ Sequence Read Archive (DRA) under the accession PRJDB40438 for scRNA-seq data of *Meiosin*-KO, *Meiosin*-ΔbHLH, and *Stra8*-null PRJDB40439 for zebrafish adult testes and ovaries. Sequence data for the mouse *Meiosin-like* gene are deposited in the National Center for Biotechnology Information-National Institutes of Health (NCBI-NIH) GenBank under accession numbers LC649921.

## Materials and Resource Availability

Mouse lines generated in this study have been deposited to Center for Animal Resources and Development (CARD ID3012-3015 for *Meiosin* ΔbHLH mutant mouse, CARD ID2999 for *Meiosin*-3xFLAG-HA knockin mouse). The antibodies are available upon request. There are restrictions to the availability of antibodies due to the lack of an external centralized repository for its distribution and our need to maintain the stock. We are glad to share antibodies with reasonable compensation by requestor for its processing and shipping.

## Acknowledgments

The authors thank Kumi Matsuura, Kazumasa Takemoto, Kie Shioya, Fabien Velilla, Miki Ohno and Yoshimi Yamazaki for technical assistance, and Koichi Kawakami for provision of zebrafish line *Tg(gata6SAGFF254A, UAS::NTR*). This work was supported in part by KAKENHI grant (20K22638, 22K15039, 25K18480) to R.S. ; Grants from The Mochida Memorial Foundation for Medical and Pharmaceutical Research; The Takeda Science Foundation; Astellas Foundation for Research on Metabolic Disorders to R.S. KAKENHI grant 25H01308 to S.K. KAKENHI grants (21KK0129, 21K06159) to N.S. KAKENHI grants (22K19315, 23H00379, 24H02050, 25H01297, 25H01304, 22H04922 AdAMS) to K.I. ; Grant from AMED PRIME (21gm6310021h0003) to K.I. ; The Mitsubishi Foundation; The Naito Foundation ; Astellas Foundation for Research on Metabolic Disorders; The Uehara Memorial Foundation ; Takeda Science Foundation to K.I ; the program of the Research for Inter-University Research Network for High Depth Omics, IMEG, Kumamoto University to S.K. and K.I.

## Author contributions

RS, KI performed most of experiments and wrote the draft of the manuscript.

YI, TK, NS performed zebrafish experiments.

SK performed phylogenetic analysis.

HN performed interaction assays.

SI generated antibodies.

SF performed histological analysis.

NT performed MS analysis.

KA generated mutant mice and performed IVF.

KY SU assisted scRNA-seq analysis

The experimental design and interpretation of data were conducted by RS, YI, NS, SK and KI.

---

**Supplementary Figure 1.**
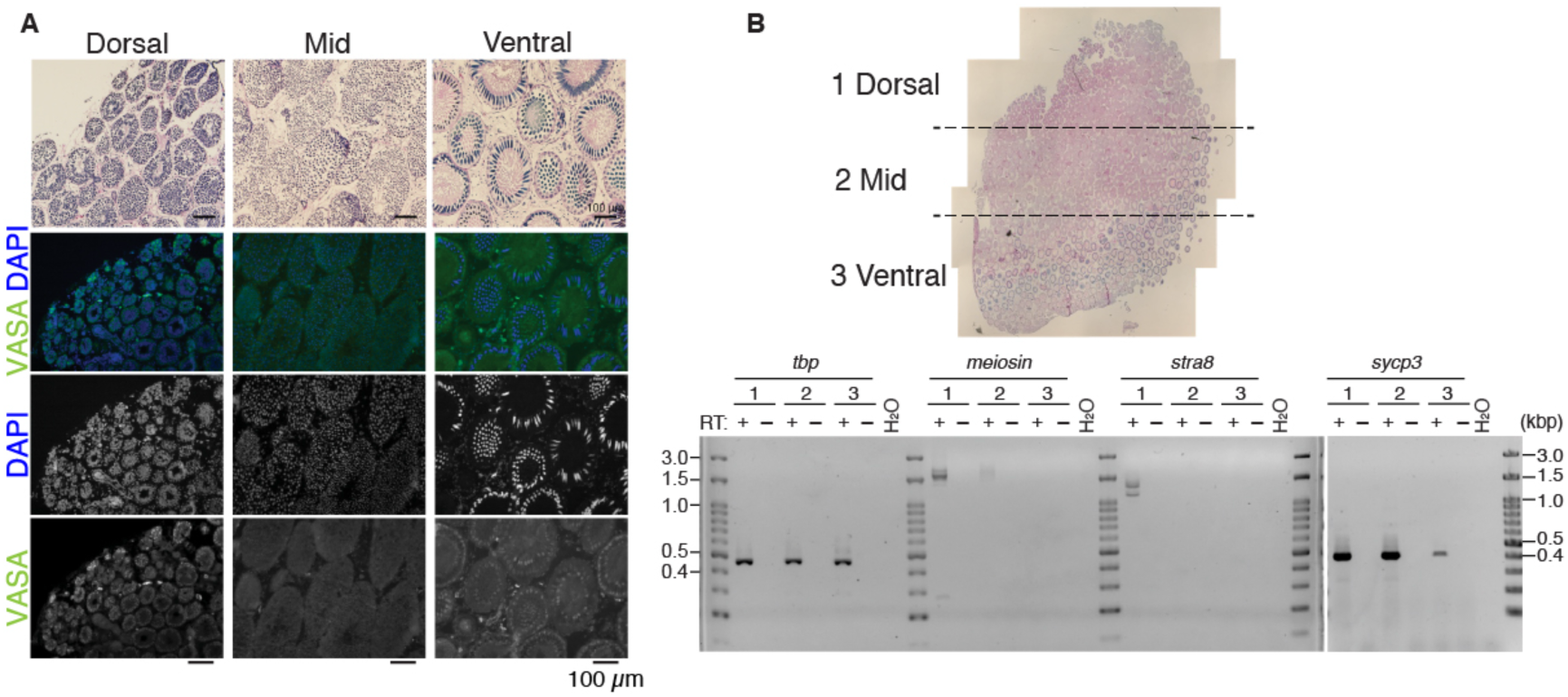
Meiosin ortholog identified in cloudy catshark (related to Figure 1) **(A)** Cloudy catshark (*Scyliorhinus torazame*) adult testis sections stained with hematoxylin and eosin (top) or immunostained as indicated (bottom). Scale bar, 100 μm. **(B)** RNA was isolated from the dorsal (1), mid (2), and ventral (3) regions of cloudy catshark testis and analyzed by RT–PCR.

**Supplementary Figure 2.**
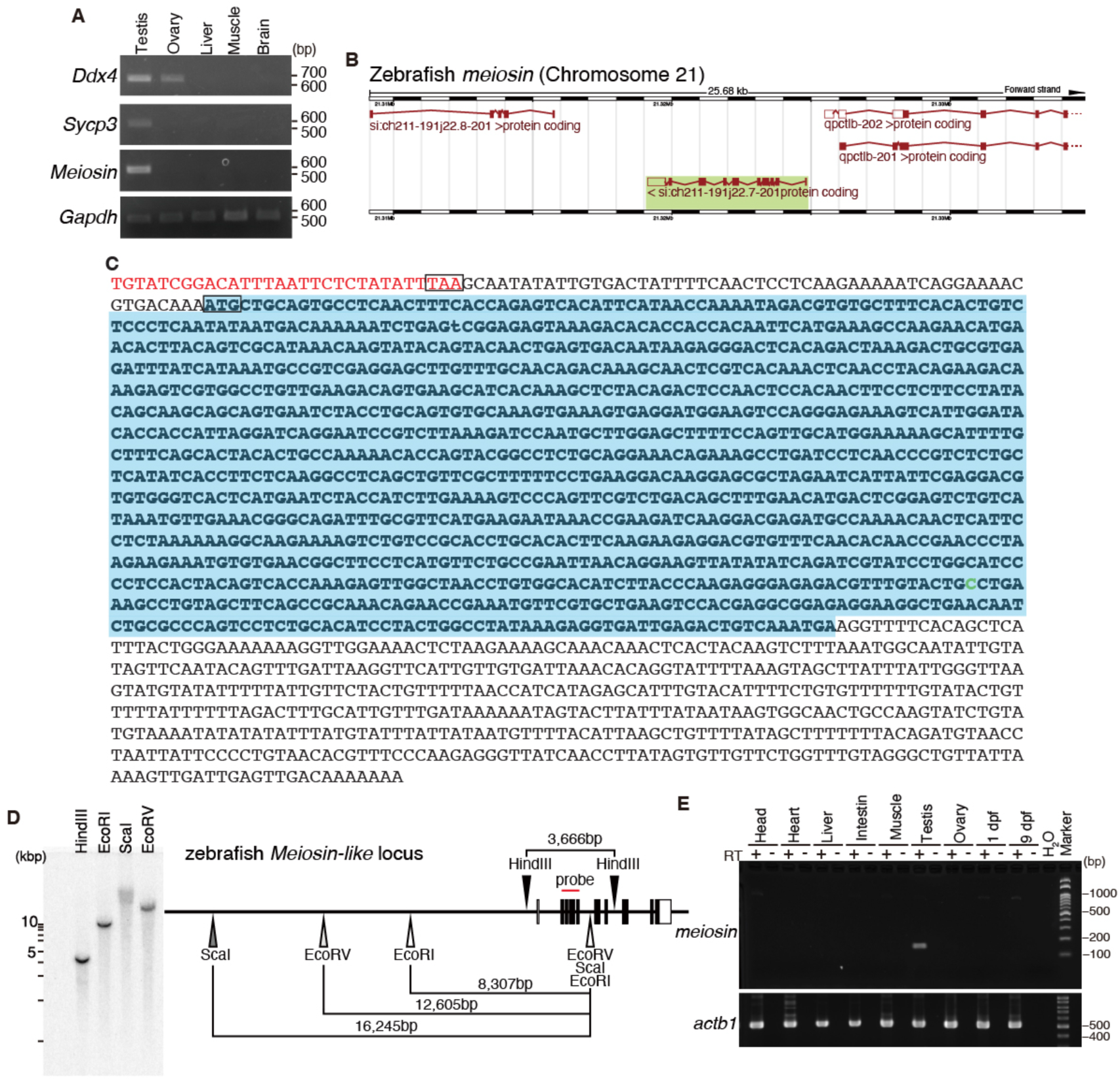
Meiosin orthologs identified in zebrafish and hagfish (related to Figure 1) **(A)** Tissue-specific expression of hagfish (*Eptatretus burgeri*) *Meiosin* examined by RT–PCR. RNA was isolated from adult male and female tissues. **(B)** Genomic view of the zebrafish (*Danio rerio*) *Meiosin* ortholog locus (si:ch211-191j22.7) on chromosome 21. **(C)** cDNA sequence of the putative zebrafish meiosin transcript derived from si:ch211-191j22.7. The blue box indicates the open reading frame. Red text marks the sequence identified by 5′ RACE. The first ATG codon and an in-frame TAA codon in the 5′ UTR are boxed. The cytosine insertion identified by cDNA sequencing that is absent from the previously annotated transcript si:ch211-191j22.7-201 is colored in green. **(D)** Southern blot analysis of zebrafish genomic DNA digested with the indicated restriction enzymes (left). A schematic of the zebrafish *Meiosin* locus, including restriction sites and probe position, is shown on the right. **(E)** Tissue-specific expression of zebrafish *Meiosin* examined by RT–PCR. RNA was isolated from adult male and female tissues and from whole embryos at 1 and 9 dpf. RT− indicates control PCR performed without reverse transcription.

**Supplementary Figure 3.**
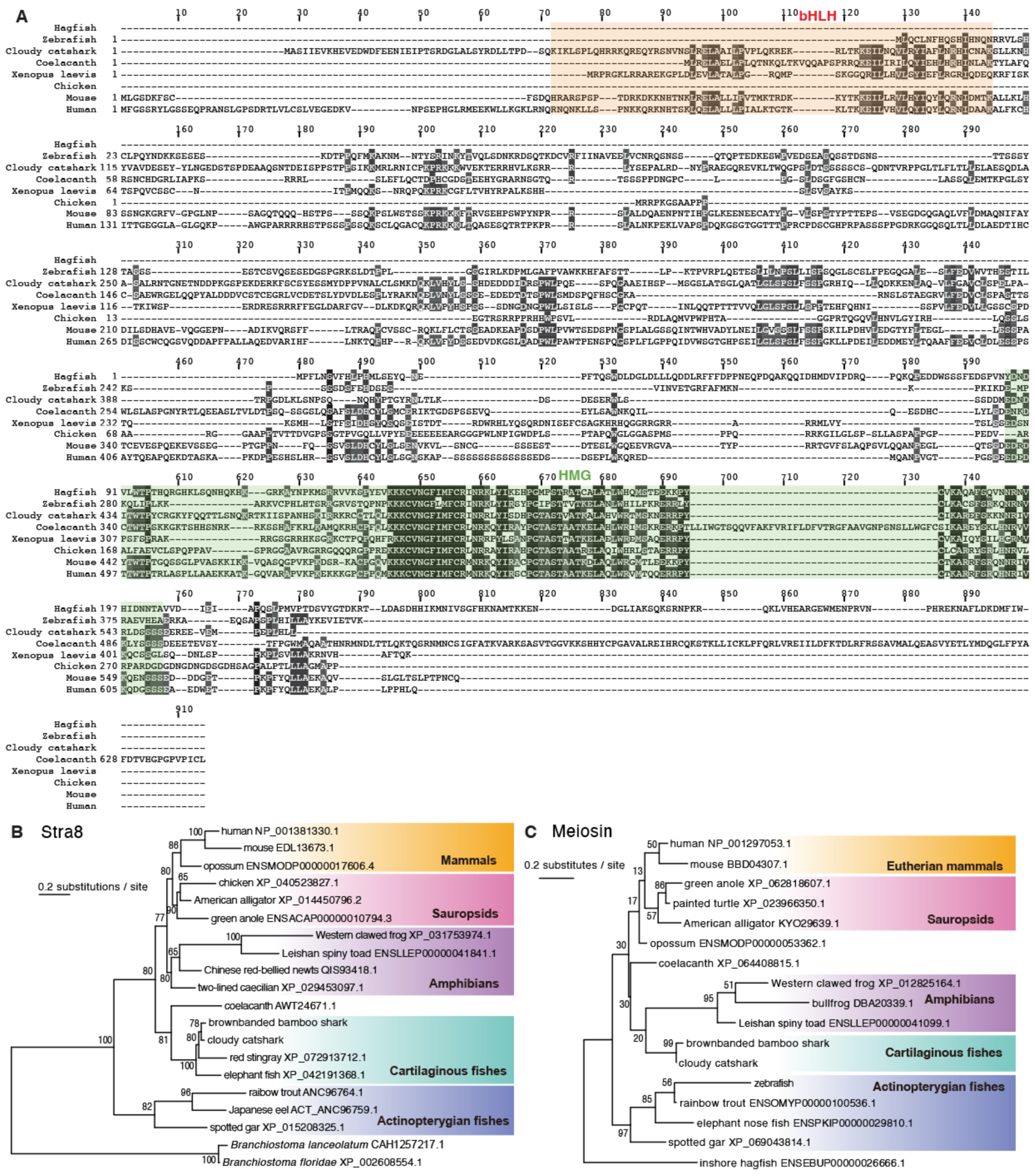
Evolutional conservation of MEIOSIN and STRA8 orthologs across vertebrates (related to Figure 1) **(A)** Sequence alignment of MEIOSIN orthologs from hagfish, zebrafish, cloudy catshark, coelacanth, Xenopus, chicken, mouse, and humans. The bHLH and HMG domains were predicted using Phyre2 based on mouse MEIOSIN and are indicated in red and green, respectively. **(B, C)** Phylogenetic trees of STRA8 (B) and MEIOSIN (C) orthologs. See Methods for methodological details.

**Supplementary Figure 4.**
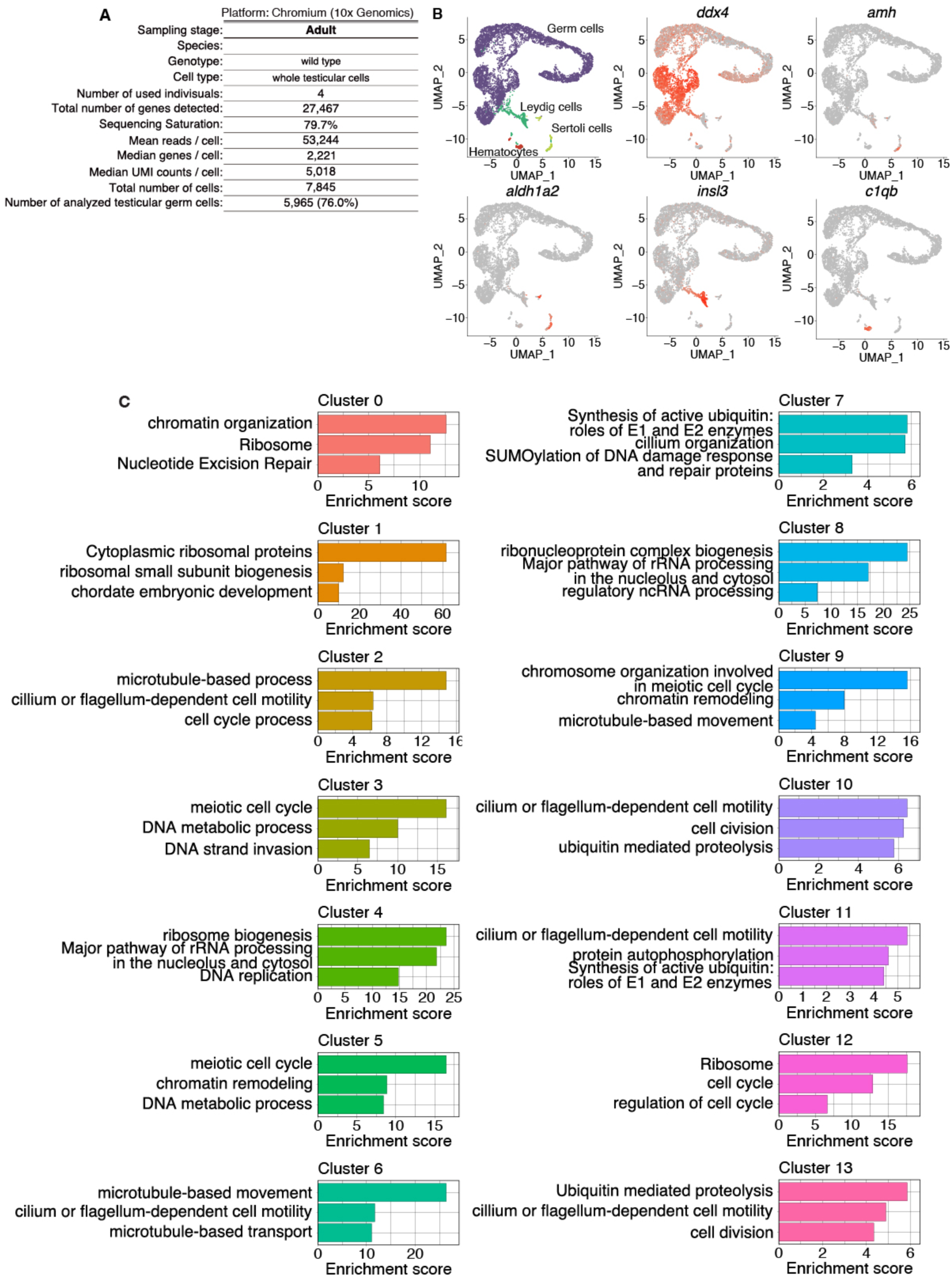
Clustering analysis of scRNA-seq dataset of whole testicular cells from adult wild type zebrafish (related to Figure 2) **(A)** Summary of 10x Genomics Chromium metrics for scRNA-seq analysis of testicular germ cells from WT adult males. The number of pooled testes and ovaries is indicated. The total number of germ cells analyzed and separated from somatic cells, as well as the percentage of germ cells among total captured cells, are shown. **(B)** UMAP representation of scRNA-seq transcriptomes from WT testicular cells. UMAP plots show mRNA abundance of key cell type-specific markers: *ddx4*, germ cells; *amh* and *aldh1a2*, Sertoli cells; *insl3*, Leydig cells; *c1qb*, hematocytes. **(C)** Gene enrichment analysis of differentially expressed genes in UMAP-defined clusters of zebrafish testicular germ cells. See also Supplementary Data 1.

**Supplementary Figure 5.**
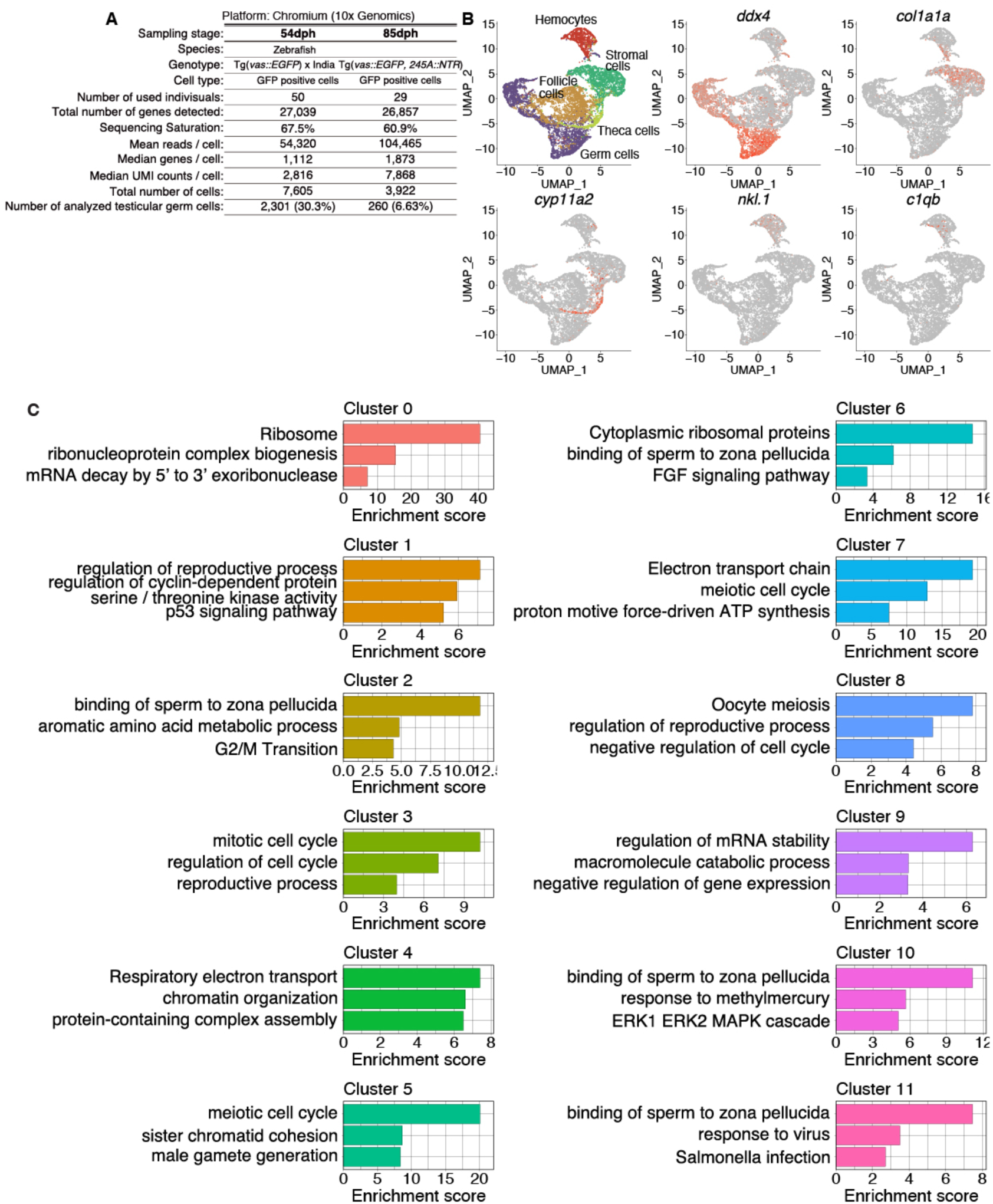
Clustering analysis of zebrafish testicular germ cells (related to Figure 2) **(A)** Summary of 10x Genomics Chromium metrics for scRNA-seq analysis of ovarian germ cells from Tg(*vas::EGFP*) females at 54 and 85 dpf. The number of pooled ovaries is indicated. The total number of germ cells analyzed and separated from somatic cells, as well as the percentage of germ cells among total captured cells, are shown. **(B)** UMAP representation of scRNA-seq transcriptomes from ovarian cells of Tg(*vas::EGFP*) females. UMAP plots show mRNA abundance of key cell type-specific markers: *ddx4*, germ cells; col1a1a, stromal cells; *cyp11a2*, theca cells; *nkl.1* and *c1qb*, hematocytes. **(C)** Gene enrichment analysis of differentially expressed genes in UMAP-defined clusters of zebrafish ovarian germ cells. See also Supplementary Data 2.

**Supplementary Figure 6.**
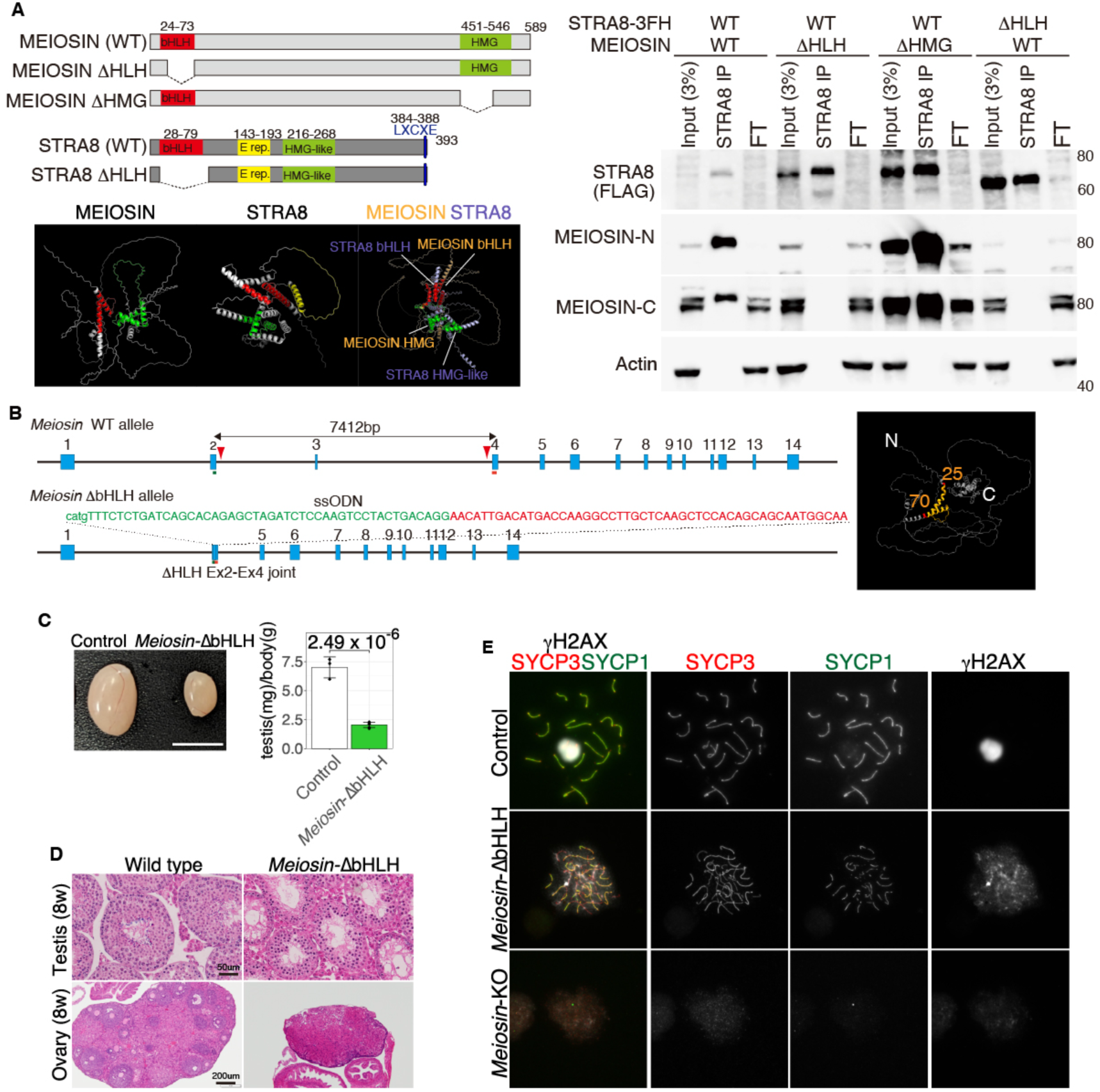
The bHLH domain of MEIOSIN is required for full meiotic function (Related to Figure 4) **(A)** Domain architecture and motifs in mouse MEIOSIN and STRA8 (upper left), and predicted protein structures generated by AlphaFold2 (lower left). Corresponding domains are color-coded as in the schematics. Right, western blot analysis of anti-FLAG immunoprecipitates from 293T cell extracts expressing the indicated combinations of WT and mutant STRA8 and MEIOSIN proteins. FT, flow-through. STRA8-3×FLAG-HA was detected with anti-FLAG antibody. MEIOSIN was detected with antibodies recognizing the N terminus or C terminus. Two independent experiments yielded similar results. **(B)** Schematic of the WT *Meiosin* allele and the *Meiosin*-ΔbHLH mutant allele. Blue boxes indicate exons. Triangle, CRISPR gRNA target site. The ssODN donor sequence is shown. The deleted region of MEIOSIN (amino acids 25–70) is indicated on the AlphaFold2-predicted structure at right. **(C)** Testes from 8-week-old control and *Meiosin*-ΔbHLH homozygous mice. Scale bar, 5 mm. Right, testis-to-body-weight ratio (mg/g) for the indicated genotypes (n = 3 each; mean ± S.D.). *** *p* = 2.94 x 10^-6^, two-sided t-test. **(D)** Hematoxylin and eosin staining of sections from 8-week-old WT and Meiosin-ΔbHLH homozygous testes (top) and ovaries (bottom). Scale bars, 50 μm (testes) and 200 μm (ovaries). **(E)** Chromosome spreads from control, *Meiosin*-ΔbHLH, and *Meiosin* KO spermatocytes at PD17 stained as indicated.

**Supplementary Figure 7.**
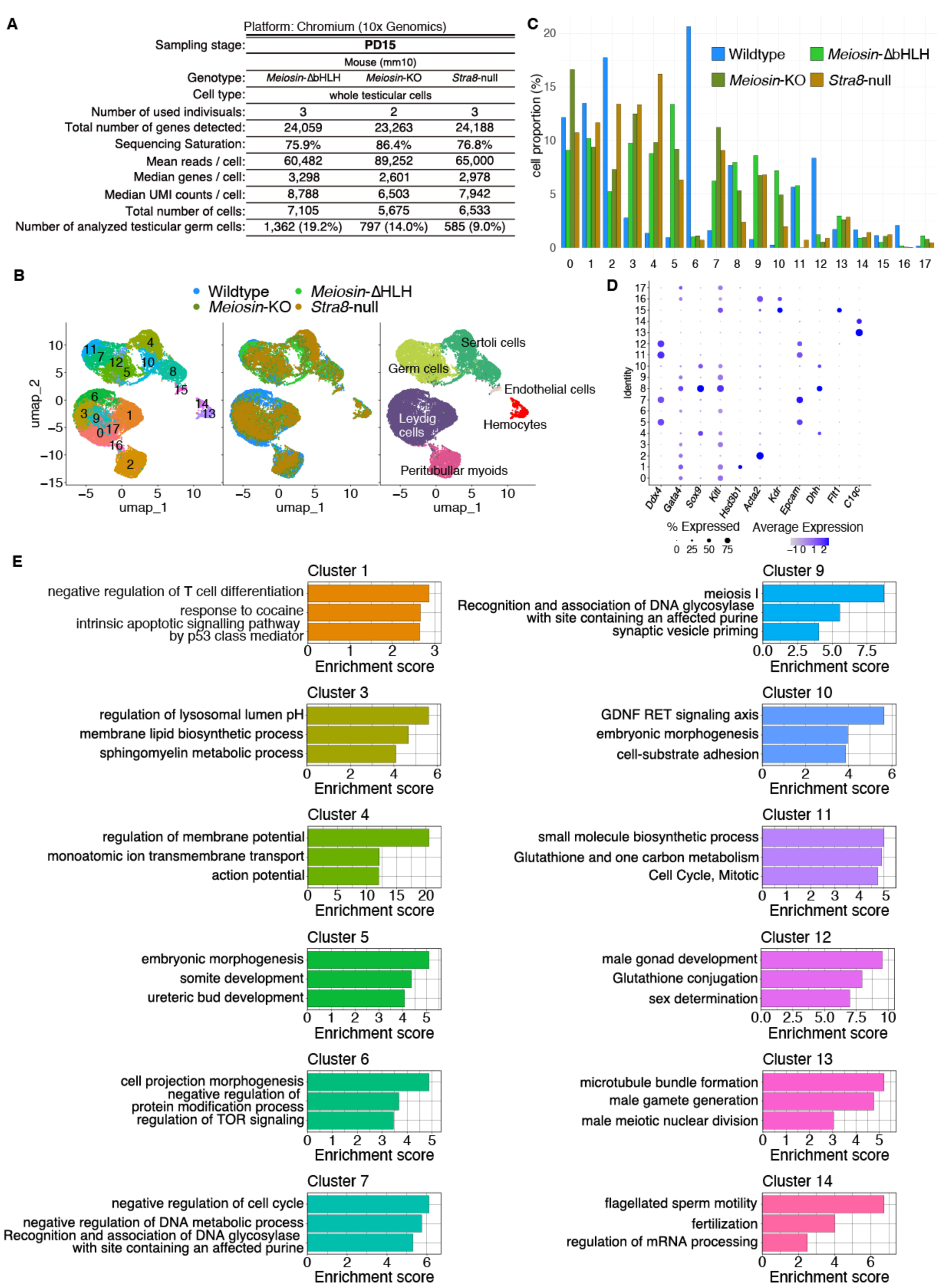
Clustering analysis of WT, *Meiosin*-ΔbHLH, *Meiosin* KO, and *Stra8*-null germ cells (Related to Figure 5) **(A)** Summary of 10x Genomics Chromium metrics for scRNA-seq analysis of testicular germ cells from *Meiosin*-ΔbHLH, *Meiosin* KO, and *Stra8*-null mice at postnatal day 15 (PD15). The number of pooled testes is indicated. Total germ cell numbers analyzed, and the percentage of germ cells among captured cells, are shown. **(B)** UMAP representation of scRNA-seq transcriptomes from testicular cells of WT, *Meiosin*-ΔbHLH, *Meiosin* KO, and *Stra8*-null mice. **(C)** Proportion of cells from each genotype in each cluster. **(D)** Dot plot showing average scaled expression (color gradient) and the fraction of cells with detectable expression (dot size) for marker genes across UMAP-defined clusters. **(E)** Metascape analysis of the top 100 enriched genes in Clusters 1, 3, 4, 5, 6, 7, 9, 10, 11, 12, 13, and 14. Bars indicate enrichment scores -log10(*p-*value).

**Supplementary Figure 8.**
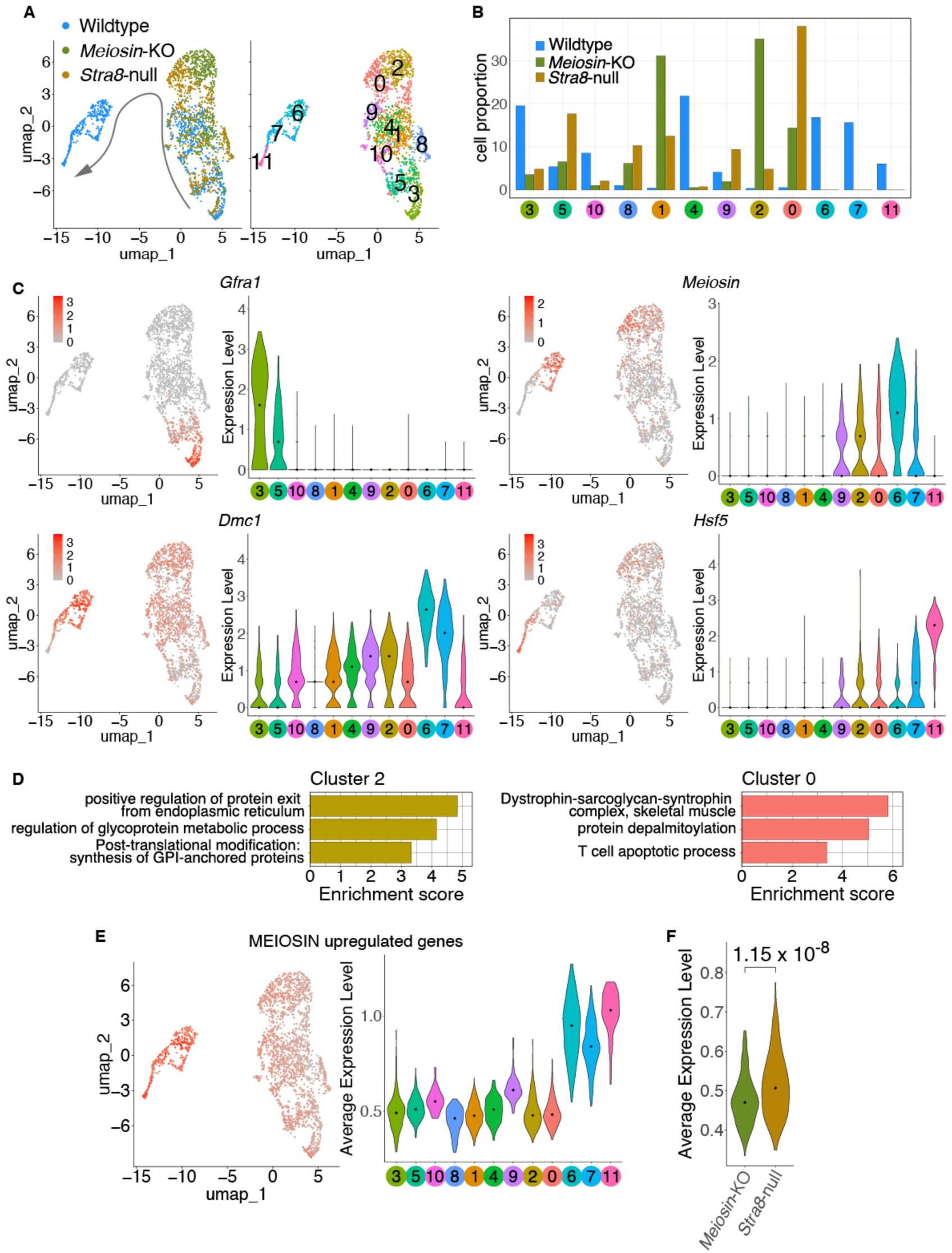
scRNA-seq analysis of WT, *Meiosin*-KO and *Stra8*-null spermatogenic germ cells (Related to Figure 6) **(A)** UMAP representation of scRNA-seq transcriptomes from spermatogenic germ cells isolated from PD15 WT, *Meiosin* KO, and *Stra8*-null testes. Cells are colored by genotype (left) or cluster identity (right). Arrow indicates the inferred differentiation trajectory based on marker gene expression. **(B)** Proportion of each genotype in each cluster. **(C)** UMAP plots (left) and violin plots (right) showing mRNA abundance of key marker genes in spermatogenic germ cells. Black dots indicate medians. Marker genes: *Gfra1*, spermatogonial stem cells; *Meiosin*, meiotic initiation; *Dmc1*, early meiotic prophase; *Hsf5*, late meiotic prophase. **(D)** Metascape analysis of genes enriched in Clusters 2 and 0. See also Supplementary Data7 for remaining clusters. **(E)** Average expression of MEIOSIN-upregulated genes shown on UMAP (left) and in violin plots (right). **(F)** Violin plots showing average expression of MEIOSIN-upregulated genes in cells assigned to Clusters 0 and 2. Statistical significance is shown by *p*-value (*p* =1.15 x 10^-8^, two-sided Wilcoxon rank sum test)

**Supplementary Figure 9.**
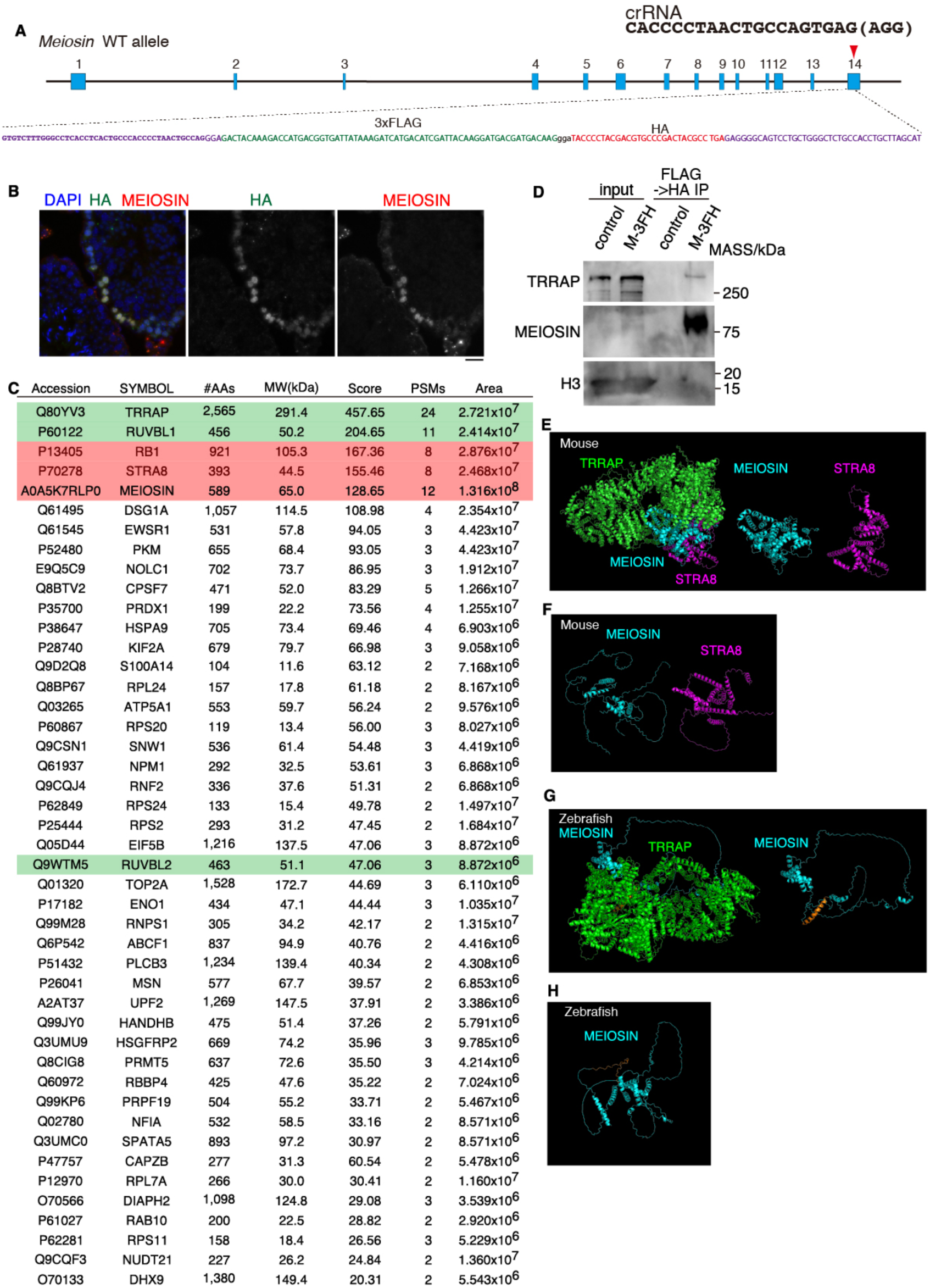
MEIOSIN interacts with components of the NuA4 histone acetyltransferase complex (Related to Figure 6) **(A)** Schematic of the Meiosin-3×FLAG-HA knock-in allele. **(B)** Seminiferous tubule sections from adult Meiosin-3FH testes stained for HA, MEIOSIN, and DAPI. Scale bar, 20 μm. **(C)** Immunoprecipitated proteins identified by more than two distinct peptide hits are listed together with their Swiss-Prot accession numbers, gene symbols, amino acid lengths, Mascot scores, and peptide hit counts. Components of the NuA4/TIP60 complex are highlighted in green. MEIOSIN, STRA8, and RB1, previously identified in STRA8 immunoprecipitates, are highlighted in red. See also Supplementary Data8. **(D)** Western blot analysis of proteins recovered by tandem affinity purification using anti-FLAG and anti-HA antibodies from chromatin extracts of non-tagged control and Meiosin-3FH knock-in testes. **(E)** Predicted 3D structure of the mouse TRRAP (green)–MEIOSIN (cyan)–STRA8 (magenta) complex (left). Ribbon models of MEIOSIN and STRA8 within the complex are shown at center and right, respectively. **(F)** Predicted 3D structures of mouse MEIOSIN and STRA8 alone. **(G)** Predicted 3D structure of the zebrafish TRRAP (green)–MEIOSIN (cyan) complex (left). The zebrafish MEIOSIN ribbon model within the complex is shown at right. The orange region denotes a segment predicted to become structured upon complex formation with TRRAP. **(H)** Predicted 3D structure of zebrafish MEIOSIN alone. The orange region corresponds to the segment predicted to gain structure upon complex formation with TRRAP.

